# 3D mapping of host-parasite-microbiome interactions reveals metabolic determinants of tissue tropism and disease tolerance in Chagas disease

**DOI:** 10.1101/727917

**Authors:** Ekram Hossain, Sharmily Khanam, Chaoyi Wu, Sharon Lostracco-Johnson, Diane Thomas, Mitchelle Katemauswa, Camil Gosmanov, Danyang Li, Christine Woelfel-Monsivais, Krithivasan Sankaranarayanan, Laura-Isobel McCall

## Abstract

Chagas disease (CD) is a parasitic infection caused by *Trypanosoma cruzi* protozoa. Over 8 million people worldwide are *T. cruzi*-positive, 20-30% of which will develop cardiomyopathy, megaoesophagus and/or megacolon. The mechanisms leading to gastrointestinal (GI) symptom development are however poorly understood. To address this issue, we systematically characterized the spatial impact of experimental *T. cruzi* infection on the microbiome and metabolome across the GI tract. The largest microbiota perturbations were observed in the proximal large intestine in both acute and chronic disease, with chronic-stage effects also observed in the cecum. Strikingly, metabolomic impact of acute-to-chronic stage transition differed depending on the organ, with persistent large-scale effects of infection primarily in the oesophagus and large intestine, providing a potential mechanism for GI pathology tropism in CD. Infection particularly affected acylcarnitine and lipid metabolism. Building on these observations, treatment of infected mice with carnitine-supplemented drinking water prevented acute-stage mortality with no changes in parasite burden. Overall, these results identified a new mechanism of disease tolerance in CD, with potential for the development of new therapeutic regimens. More broadly, these results highlight the potential of spatially-resolved metabolomic approaches to provide insight into disease pathogenesis, with translational applications for infectious disease drug development.

## Introduction

Chagas disease (CD), also known as American trypanosomiasis, is a neglected tropical disease endemic in Latin America ^1^. However, due to migration CD now has a global reach spanning North America, Europe and Asia ^2^. Six to eight million people are infected with *T. cruzi*, with approximately 12,000 deaths per year ^3^. CD is caused by infection with the protozoan parasite *Trypanosoma cruzi*. Infected individuals pass first through an acute disease stage, usually asymptomatic, then to a chronic asymptomatic (indeterminate) stage that can last for decades. Thirty to forty percent of infected individuals progress from indeterminate to determinate (symptomatic) chronic CD ^4^, 20-30% with cardiovascular complications (heart failure, arrhythmias, and thromboembolism) and 15-20% of infected individuals with gastrointestinal (GI) symptoms (megaesophagus and megacolon) ^5^. Digestive CD has been neglected compared to cardiac CD and consequently is much more poorly understood. However, recent studies using bioluminescent parasites in mouse models have shown that specific sites in the GI tract are parasite reservoirs in chronic CD and may be major contributors to cardiac symptom development, particularly after treatment failure ^6,7,8^. Treatment of GI CD is also challenging, with limited options once symptoms become apparent ^1^. There is therefore a strong need to improve our understanding of the interaction between *T. cruzi* and the GI tract, both to clarify mechanisms of GI CD pathogenesis, and to define GI factors contributing to cardiac CD, leading to new treatment strategies.

The GI tract is a complex environment where host, pathogen and microbiota interact to affect disease pathogenesis ^9^. We previously demonstrated that *T. cruzi* infection affects the fecal microbiome and metabolome, but information on the specific GI sites driving this output had not yet been determined ^10^. In this study, we applied a novel integration of small molecule-focused liquid chromatography-tandem mass spectrometry (LC-MS/MS) and 3D modeling (“chemical cartography”), in conjunction with microbiome analysis, to systematically characterize the *T. cruzi-*induced changes in the GI microenvironment in acute and chronic CD. We specifically focused on small molecule characterization because they represent the output of cellular processes as well as their regulators, and therefore have the closest relationship to phenotype ^11^. Given that most drugs are still small molecule-based ^12^, we further hypothesized that identifying infection-associated disturbances in the small molecule profile can most rapidly lead to new treatments for CD.

Results identified organ-specific and organ sub-site-specific magnitudes of disruptions in the chemical and microbial GI environment by *T. cruzi*, and highlighted differential mechanisms of acute to chronic stage transitions depending on organ. These results provide a mechanism by which consistent perturbations of tissue biochemical pathways lead to GI CD pathology in the oesophagus and large intestine. Consistent infection-induced elevation of acylcarnitine family members across organs further led us to investigate the role of acylcarnitines in disease pathogenesis. Supplementing animal drinking water with carnitine prevented acute-stage mortality in experimental CD in the absence of antiparasitic effect, revealing a novel mechanism of disease tolerance in CD. Overall, these results identified novel mechanisms of CD pathogenesis, with major translational applications to CD drug development. Furthermore, the data collected here on uninfected animals, and our approach in general, can serve as a reference to investigate determinants of tropism and novel treatment strategies for any other GI pathogen.

## Results

### Regiospecific molecular impact of *T. cruzi* colonization in the GI tract

GI CD is still poorly understood. In our prior work, we identified specific small molecules correlated with cardiac parasite tropism ^13^. Here, we sought to identify the locoregional chemical changes associated with parasite GI colonization. Mice were infected with 1,000 luciferase-expressing *T. cruzi* strain CL Brener parasites ^6^. Twelve days (acute stage) and 89 days (chronic stage) post-infection, animals were euthanized, the GI tract sectioned (**Fig. S1a**), and parasite burden in each section determined by *ex vivo* bioluminescence imaging (**Fig. 1abc**). At 12 days post infection, parasite burden was high throughout the GI tract, with the highest parasite burden in the distal small intestine (position 9; p<0.05 Student’s T-test, distal small intestine to mid and distal stomach, small intestine position 6 and small intestine position 7), and the lowest parasite burden in the cecum (position 10, p<0.05 Student’s T test cecum vs oesophagus, distal small intestine and distal large intestine; **Fig. 1ab)**. In contrast, at 89 days post-infection, the parasite burden was highest in the cecum (p<0.05 Student’s T-test, cecum vs oesophagus, proximal and distal small intestine), and lowest in the proximal small intestine (p<0.05 Student’s T-test, small intestine positions 5 and 6 vs oesophagus, proximal and distal large intestine, **Fig. 1ac**). In general, as expected, parasite burden decreased from the acute stage to the chronic stage (**Fig. S1b**). However, surprisingly, parasite burden increased in the cecum during the acute to chronic transition, suggesting a possible role for the cecum as a parasite reservoir protected from antiparasitic immune responses (**Fig. S1b**). These observations support the concept that all GI sites can initially harbor *T. cruzi*, but then differentially respond to parasite presence, leading to the ability of the parasite to persist and cause damage in some sites but not others.

**Fig. 1.**
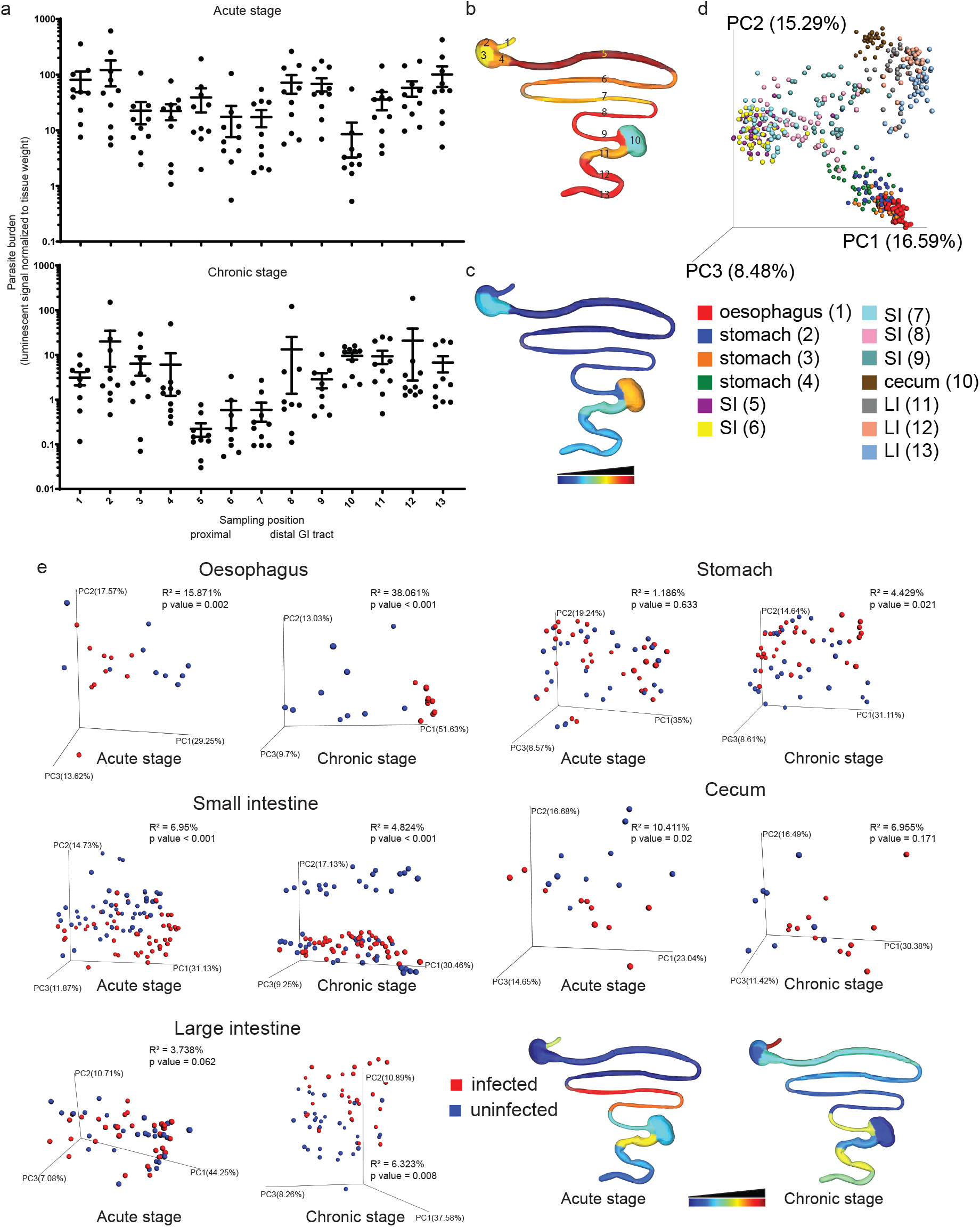
Spatial impact of *T. cruzi* infection is reflected by spatial modulation of the tissue small molecule profile. C3H/HeJ male mice (n=5 per group and per replicate) were mock-infected or infected with 1,000 luminescent *T. cruzi* strain CL Brener trypomastigotes, in two biological replicates. GI samples were collected 12 and 89 days post-infection. (a) Parasite burden at each sampling site. GI was sectioned into 13 segments, and luminescence quantified at each site. To correct for variations in sample size, luminescence counts were normalized to sample weight, for each sample. Mean + standard error of mean are displayed. (b) Median luminescent signal at each sampling site, 12 days post-infection. Sampling positions are indicated. (c) Median luminescent signal at each sampling site, 89 days post-infection. Common scale for b and c. (d) Principal coordinate analysis showing separation between sampling sites in terms of overall chemical composition, even within a given organ (negative mode, all timepoints combined, Bray-Curtis-Faith distance metric; p<0.001 R2=13.918%). (e) Principal coordinate analysis showing significant chemical composition differences between infected and uninfected samples in oesophagus (acute stage, PERMANOVA p=0.002, R^2^=15.871%; chronic stage PERMANOVA p<0.001, R^2^=38.061%), stomach (chronic stage, PERMANOVA p=0.021, R^2^=4.429%; non-significant acute stage), small intestine, (acute stage, PERMANOVA p<0.001, R^2^=6.95%; chronic stage, PERMANOVA p<0.001, R^2^=4.824), cecum (acute stage, PERMANOVA p=0.02, R^2^=10.411%; non-significant chronic stage) and large intestine (chronic stage, PERMANOVA p=0.008, R^2^=6.323%; non-significant acute stage). Bottom right-hand panels display R^2^ at each sampling site (common logarithmic scale, acute and chronic stage).

Given this differential parasite tropism in the chronic stage and the unique aspects of CD pathology, we sought to investigate the molecular determinants of parasite persistence vs disease resolution. To do so, we extracted small molecules (metabolites) from each collected GI section (637 samples total) and analyzed these molecules by LC-MS/MS in positive and in negative mode. As expected, the strongest determinant of overall chemical profile was the source organ, and sample position within that organ, as observed by principal coordinate (PCoA) analysis (**Fig. 1d**, PERMANOVA based on source organ, p<0.001, R^2^=46.374% for positive mode LC-MS/MS analysis and p<0.001, R^2^=47.033 % for negative mode LC-MS/MS analysis; PERMANOVA based on sampling position, p<0.001, R^2^=13.494% for positive mode LC-MS/MS analysis and p<0.001, R^2^=16.477% for negative mode LC-MS/MS analysis (12 days post-infection); PERMANOVA based on sampling position, p<0.001 R^2^=12.067% positive mode and p<0.001 R^2^=13.918% negative mode (all timepoints combined)). Overall impact of infection was much more minor (**Fig. S2, S3**, PERMANOVA based on infection status, acute stage, p=0.019, R^2^=1.011% for positive mode LC-MS/MS analysis and p=0.07, R^2^=0.692% for negative mode LC-MS/MS analysis; PERMANOVA based on infection status, chronic stage, p=0.014, R^2^=0.982% for positive mode LC-MS/MS analysis and p=0.007, R^2^=0.981% for negative mode LC-MS/MS analysis). Comparison of chemical families differentially-modulated by infection also identified few commonalities between sample sites (**Fig. S4, S5**). We therefore focused our analysis on the impact of infection in each individual organ. Visualization of the chemical profile in each organ in relationship to infection status using PCoA analysis revealed organ-specific differences in the impact of *T. cruzi* infection. Acute-stage infection was associated with major disturbances in the overall oesophagus chemical profile (PERMANOVA p=0.002, R^2^=15.871%), with lower-scale perturbations in the small intestine and cecum (PERMANOVA p<0.001, R^2^=6.95% and PERMANOVA p=0.02, R^2^=10.411%, respectively) (**Fig. 1e**, **S6**). The strongest acute-stage disruption within the small intestine chemical environment was observed in the distal small intestine, where parasite burden is the highest (**Fig. 1e**, PERMANOVA p<0.001, R^2^=30.198%; p<0.001, R^2^=26.063; p=0.006, R^2^=12.564, for positions 7, 8, 9, respectively). These changes resolved in the chronic stage for the cecum (PERMANOVA p=0.171, R^2^=6.955%), decreased in magnitude for the small intestine (PERMANOVA p<0.001, R^2^=4.824), became apparent in the stomach and large intestine (PERMANOVA p=0.021, R^2^=4.429% and PERMANOVA p=0.008, R^2^=6.323%, respectively), and increased in magnitude in the oesophagus (PERMANOVA p<0.001, R^2^=38.061%). Importantly, the largest statistically significant sites of metabolome disturbance in the chronic stage were the oesophagus and large intestine, which are the sites of damage in symptomatic chronic-stage GI CD ^14^. On a per-sampling site basis, spatial heterogeneity in terms of overall effect size (R^2^) was observed within a given organ. The largest increase in R^2^ during the transition from the acute to the chronic stage were observed in the oesophagus, distal stomach and central large intestine (2.4, 2.0 and 1.9-fold increases, respectively).

Next, we investigated the nature of the chemical shifts associated with these infection-altered chemical profiles. Feature annotation rates were considerably higher in positive mode than in negative mode (35.4% vs 10.2%), so we focused this analysis on our positive mode LC-MS/MS data. We used machine learning (random forest) approaches to identify specific molecular features driving the differences between infected and uninfected tissues. Given our observations on the impact of sampling position on metabolite features (PCoA, **Fig. 1d**, **Fig. S2, Fig. S3, Fig. S5**), these comparisons were independently performed for each organ. In the acute stage, we observed elevation in specific acylcarnitines and specific phosphatidylcholine (PC) family members in the different organs (**Fig. 2a-f**,**Table S1, Fig. S7**). We also observed elevation in kynurenine in the stomach and large intestine in the acute stage. These differences persisted in the chronic stage for the proximal and central large intestine only (**Fig. 2g-h**,**Table S1, Fig. S7**). Strikingly, the levels of tryptophan, the precursor of kynurenine, were correspondingly decreased in the acute stage at the same large intestine sites where kynurenine was elevated, whereas it was increased by infection in the chronic stage in the oesophagus (**Fig. 2i-j**). Kynurenine is induced by inflammation; kynurenine metabolites have direct antiparasitic effects and contribute to the control of acute *T. cruzi* infection ^15^. However, they can also induce regulatory T cells ^16^, and as such, our observation of kynurenine persistence in the large intestine may contribute to parasite persistence in this organ. In accordance with our prior observations in the context of the fecal metabolome ^10^, specific large intestine and small intestine bile acid derivatives were increased in infected mice (**Fig. 2k-m**, **Table S1, Fig. S7**). At 89 days post-infection, molecular features identified as elevated by infection include specific acylcarnitines (*e.g.* C20:4 acylcarnitine in the oesophagus), specific PCs (*e.g.* PC(22:5), PC(20:4), PC(22:6) in the oesophagus; PC(22:4) in the large intestine), specific amino acids and derivatives (*e.g.* kynurenine in the large intestine, tryptophan in the oesophagus) (**Fig. 2a-j**, **Table S1, Fig. S7**). Importantly, the pattern of persistence of these metabolic changes reflected known sites of CD: for example, most of the top 10 metabolic perturbations observed in the acute stage in the oesophagus were still perturbed by infection in the chronic stage, whereas none of the small intestine acute-stage perturbations persisted in the chronic stage (**Fig. 2**, **Table S1**). Overall, these results identified tissue metabolic changes linked to CD tropism and pathogenesis, at the scale of overall chemical disturbances, as well as several metabolic pathways correlated with infection status.

**Fig. 2.**
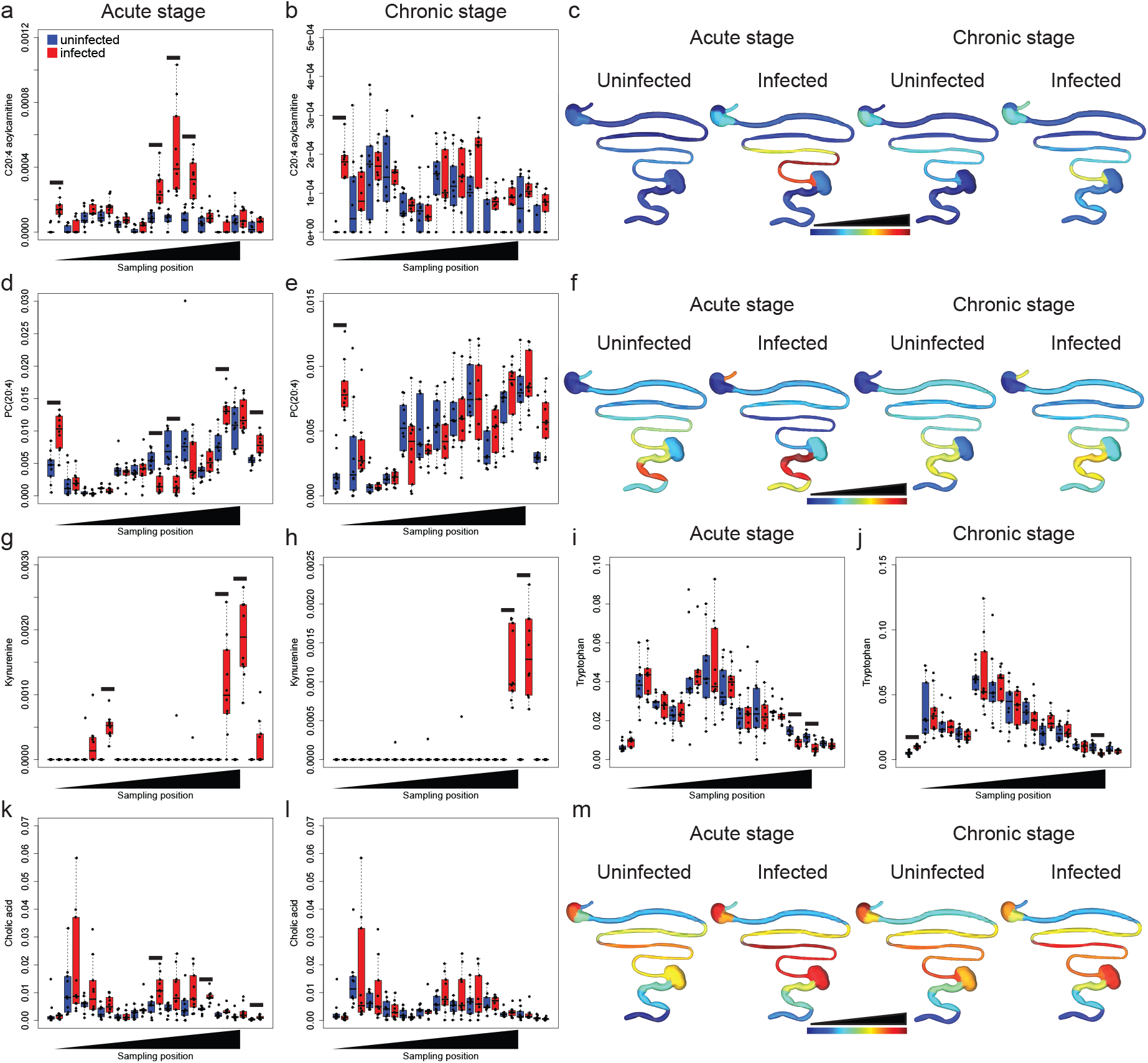
Common and tissue-specific metabolic changes identified by random forest demonstrate persistence of these alterations at sites of CD. (a) and (b) Infection-induced elevation of C20:4 acylcarnitine in the oesophagus and small intestine in the acute stage, persisting in the oesophagus in the chronic stage (FDR-corrected Mann-Whitney p=0.000988, p=0.000141, p=0.00107, p=0.00107 for acute-stage positions 1, 7, 8 and 9; p=0.0188 for chronic-stage position 1). (c) Spatial distribution of C20:4 acylcarnitine (median, common linear scale). (d) and (e) Infection-induced elevation of PC(20:4) in the oesophagus and large intestine in the acute stage, persisting in the oesophagus in the chronic stage. PC(20:4) was decreased in the acute stage in the infected small intestine (FDR-corrected Mann-Whitney p=0.00189, p=0.000844, p=0.00158, p=0.000844, p=0.000891 for acute-stage positions 1, 7, 8, 11 and 13; p=0.000141 for chronic-stage position 1). (f) Spatial distribution of PC(20:4) (median, common linear scale). (g) and (h) Infection-induced elevation of kynurenine in the stomach and large intestine in the acute stage persists in the large intestine in the chronic stage (FDR-corrected Mann-Whitney p=0.000796, p=0.00100, p=0.000796 for acute-stage positions 4, 11 and 12; p=0.000415 and p=0.000415 for chronic-stage positions 11 and 12. (i) and (j) Infection-induced decrease in tryptophan at sites of increased kynurenine in the large intestine (acute and chronic stage). Infection also increased tryptophan in the oesophagus in the chronic stage. (FDR-corrected Mann-Whitney p=0.001689 and p=0.0136 for acute-stage positions 11 and 12; p=0.000281 and p=0.0253 for chronic-stage positions 1 and 12). All detected tryptophan adducts combined. (k) and (l) Infection-induced increase in small intestine, cecum and large intestine cholic acid (all detected adducts combined), acute stage only. (FDR-corrected Mann-Whitney p=0.0498, p=0.000281, p=0.0297 for acute-stage positions 7, 10 and 13). (m) Spatial distribution of cholic acid (median, common logarithmic scale). Black lines in panels (a-e) and (g-l) indicate FDR-corrected Mann-Whitney p<0.05.

### Impact of *T. cruzi* colonization on the GI tract microbiome

Several of the molecules identified in our dataset are of microbial origin or microbially-modified, such as indole-L-lactate, indoxyl sulfate and secondary bile acids (**Fig. S8, Table S2**). Dataset match analysis through the GNPS platform ^17^ identified 1689 unique matches in our positive mode dataset to pure bacterial culture datasets not shared with pure mammalian culture datasets, known plastics-derived contaminants or blank files, suggesting a potential bacterial origin. Studies comparing germ-free and colonized mice have also shown that the microbiota influences a variety of the metabolites detected in our study, including tryptophan, tyrosine and maltotriose ^18,19^. Tryptophan and tyrosine in particular were affected by infection (tryptophan: decreased in the large intestine overall, Mann-Whitney p=6.793e-06 (acute stage) and p=0.004313 (chronic stage); tyrosine: decreased in the large intestine overall, Mann-Whitney p=0.01518 (acute stage), non-significant (chronic stage)). We have previously demonstrated that experimental *T. cruzi* infection alters the fecal microbiome and metabolome ^10^, a finding that was recently confirmed in *T. cruzi-*infected children in Bolivia ^20^. We therefore sought to evaluate the spatial impact of *T. cruzi* infection on the microbiota at each collection site in acute-stage disease (except for the oesophagus where insufficient material was available to perform both metabolomic and 16S analyses), and focusing on the cecum and large intestine in chronic disease, given their role as major sites of CD pathogenesis and the unique metabolomic pattern observed at these sites (**Fig. 1e**).

Differences in the overall microbiota composition (beta-diversity, all sites combined for a given organ) were observed in the stomach and large intestine in the acute stage (PERMANOVA p=0.05, R^2^=3.22% and PERMANOVA p=0.04, R^2^=3.837%, respectively), with non-significant changes in the small intestine and cecum (PERMANOVA p=0.069, R^2^=1.791% and PERMANOVA p=0.058, R^2^=10.657%, respectively). These differences increased in magnitude for the cecum and large intestine during the transition from acute to chronic stage (PERMANOVA p=0.02, R^2^=11.556% and PERMANOVA p=0.002, R^2^=5.83%, respectively) (**Fig. 3a-f**). Spatial heterogeneity was also observed within an organ (**Fig. 3g**), with the highest disturbances in the microbiota found in the proximal large intestine (sampling position 11, PERMANOVA p=0.022, R^2^=12.36% and PERMANOVA p=0.009, R^2^=10.715% for acute and chronic stage, respectively). Persistent disturbances in the large intestine microbiota reflect our findings for the large intestine metabolome, while the discrepancies between cecal microbiota and metabolome findings may reflect persistent luminal rather than tissue alterations. Overall, the persistence of microbiota alterations in these sites correlates well with our observation of continued alterations of the fecal microbiota and metabolome through acute and chronic experimental CD ^10^. In accordance with prior reports ^20,10^, no significant differences in alpha-diversity were observed (**Fig. S9**) between infected and uninfected tissue in both acute and chronic stages. Notably, the effect size observed for microbiota composition analysis was lower than for our tissue metabolomics analysis (**Fig. 1**), although in both cases the proximal large intestine was one of the major sites of infection-associated perturbation. This may reflect segregation of the microbiome from the site of infection, so that only indirect effects can be observed. Furthermore, cage and batch effects were found to have a larger impact on microbiome composition (**Fig. S10**), while metabolome analysis was more robust to such effects, as we previously reported ^10^.

**Fig. 3.**
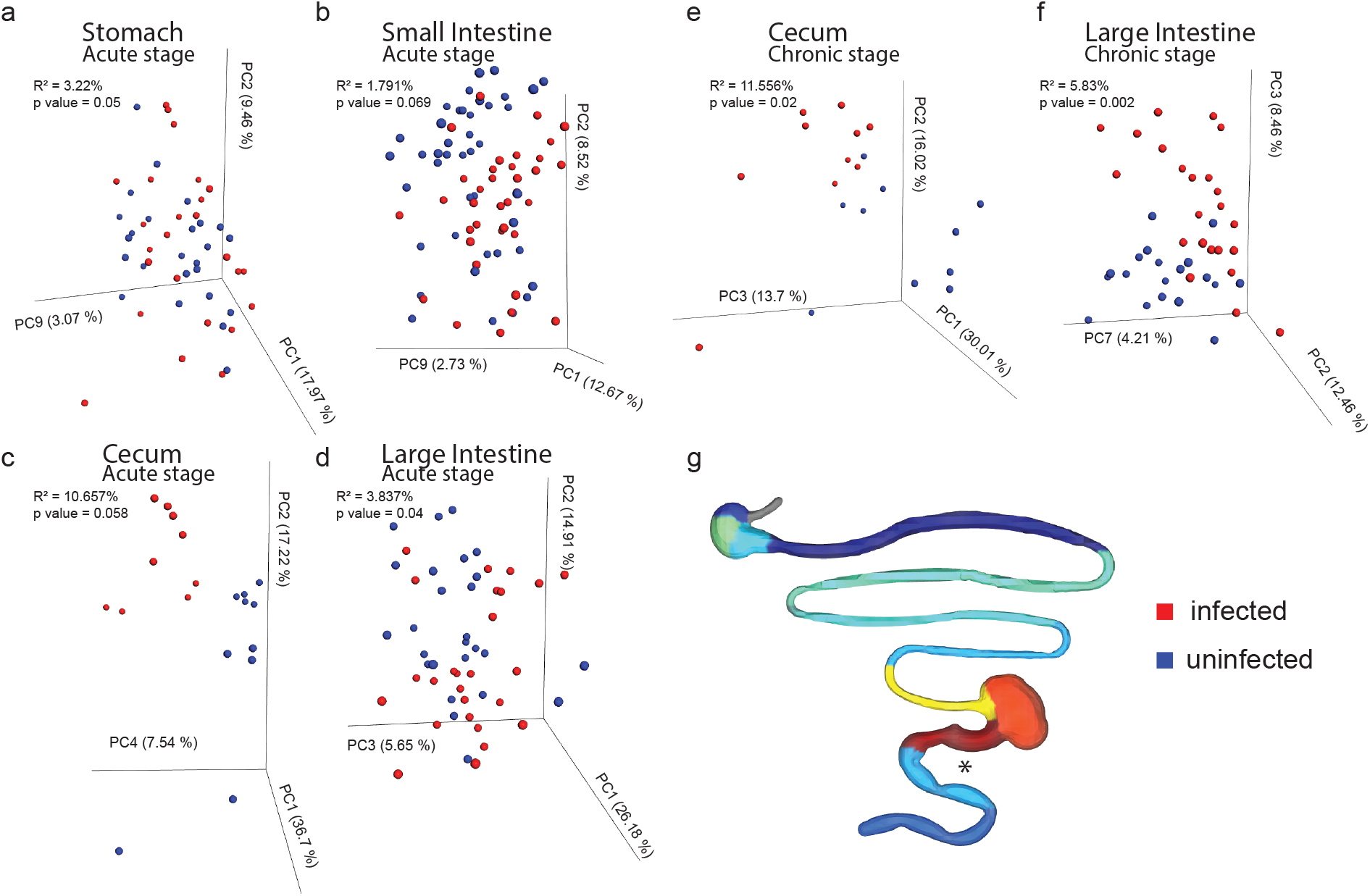
Persistent, spatially heterogeneous impact of *T. cruzi* infection on the microbiota. 16S sequencing was performed on homogenate from all sampling sites except oesophagus in the acute stage, and focusing on large intestine and cecum in the chronic stage. Principal coordinate analysis revealed significant differences in the overall microbiota composition in the stomach (a) and large intestine (d) in the acute stage (PERMANOVA p=0.05, R^2^=3.22% and PERMANOVA p=0.04, R^2^=3.837%, respectively), which persisted in the chronic stage for the large intestine (f, PERMANOVA p=0.002, R^2^=5.83%). Spatial heterogeneity was also observed within an organ (g), with the highest disturbances in the microbiota in the proximal large intestine (sampling position 11, PERMANOVA p=0.022, R^2^=12.36% and PERMANOVA p=0.009, R^2^=10.715% for acute and chronic stage, respectively). (g) displays R^2^ at each sampling site in the acute stage (logarithmic scale). *, PERMANOVA p<0.05.

### Role of acylcarnitines in CD tolerance

Translating ‘omics findings into novel therapeutic approaches is one of the major challenges of this post-genome era. Because we observed larger metabolome than microbiome infection-associated perturbations, and based on our current observations of infection-induced elevation in specific acylcarnitine family members, and our prior findings of differential cardiac acylcarnitine distribution and mass range in mild vs severe acute *T. cruzi* infection ^13^, we focused here on acylcarnitines and the potential of acylcarnitine modulation for CD treatment. The acylcarnitine sub-network (**Fig. 4a**) was manually annotated (**Table S3, Fig. S11a**), and impacts of infection on short-chain (C3-C4), mid-chain (C5-C11) and long-chain (C12 and greater) acylcarnitines assessed. While total and short-chain GI short-chain acylcarnitine levels were comparable between infected and uninfected animals 12 days post-infection (**Fig. S11bc**), we observed significant elevation in total and short-chain acylcarnitine levels at each small intestine site, in infected animals (total acylcarnitines: FDR-corrected Mann-Whitney p=0.00543, p=0.000422, p=7.04e-05, p=7.04e-05, p=0.000422 for positions 5, 6, 7, 8, 9; short-chain acylcarnitines: FDR-corrected Mann-Whitney p=0.000668, p=0.000563, p=0.000141, p=0.000563, p=0.00189 for positions 5, 6, 7, 8, 9; **Fig. S11de**). For long-chain acylcarnitines, infection-induced acylcarnitine elevation was restricted to the distal portions of the small intestine (FDR-corrected Mann-Whitney p=0.000141, p=0.000563, p=0.000563 for positions 7, 8, 9; **Fig. 4bc**). This difference between total and spatially-resolved short-chain acylcarnitine levels highlight the strength of our chemical cartography approach. Mid-chain (C5 to C11) acylcarnitine levels were not significantly different 12 days post-infection between infected and uninfected animals at any GI site (**Fig. S11f**). Acylcarnitine small intestine elevation was no longer observed in the chronic stage, except for short-chain acylcarnitines in the duodenum (sampling position 5, FDR-corrected Mann-Whitney p=0.0253), although select other GI sites showed infection-induced increases in acylcarnitines (**Fig. S11ghi**; distal large intestine, total acylcarnitines and short-chain acylcarnitines, FDR-corrected Mann-Whitney p=0.0196 and p=0.0387, respectively; oesophagus, short-chain acylcarnitines, FDR-corrected Mann-Whitney p=0.00422). Acetyl-carnitine was also elevated in select sites 12 days post-infection (FDR-corrected Mann-Whitney p=0.0.0220, p=0.003413, p=0.000141, p=0.000141, p=0.0008913, p=0.0220 for sites number 4-9 (stomach and small intestine, 12 days post-infection)), but was comparable between infected and uninfected tissues at all sites 89 days post-infection (**Fig. S11jk**). In contrast, unmodified carnitine levels were comparable throughout the intestine 12 days post-infection, and only significantly elevated in infected oesophagus and uninfected central large intestine 89 days post-infection (FDR-corrected Mann-Whitney p=0.000141 and p=0.0187, respectively) (**Fig. S11l**).

**Fig. 4.**
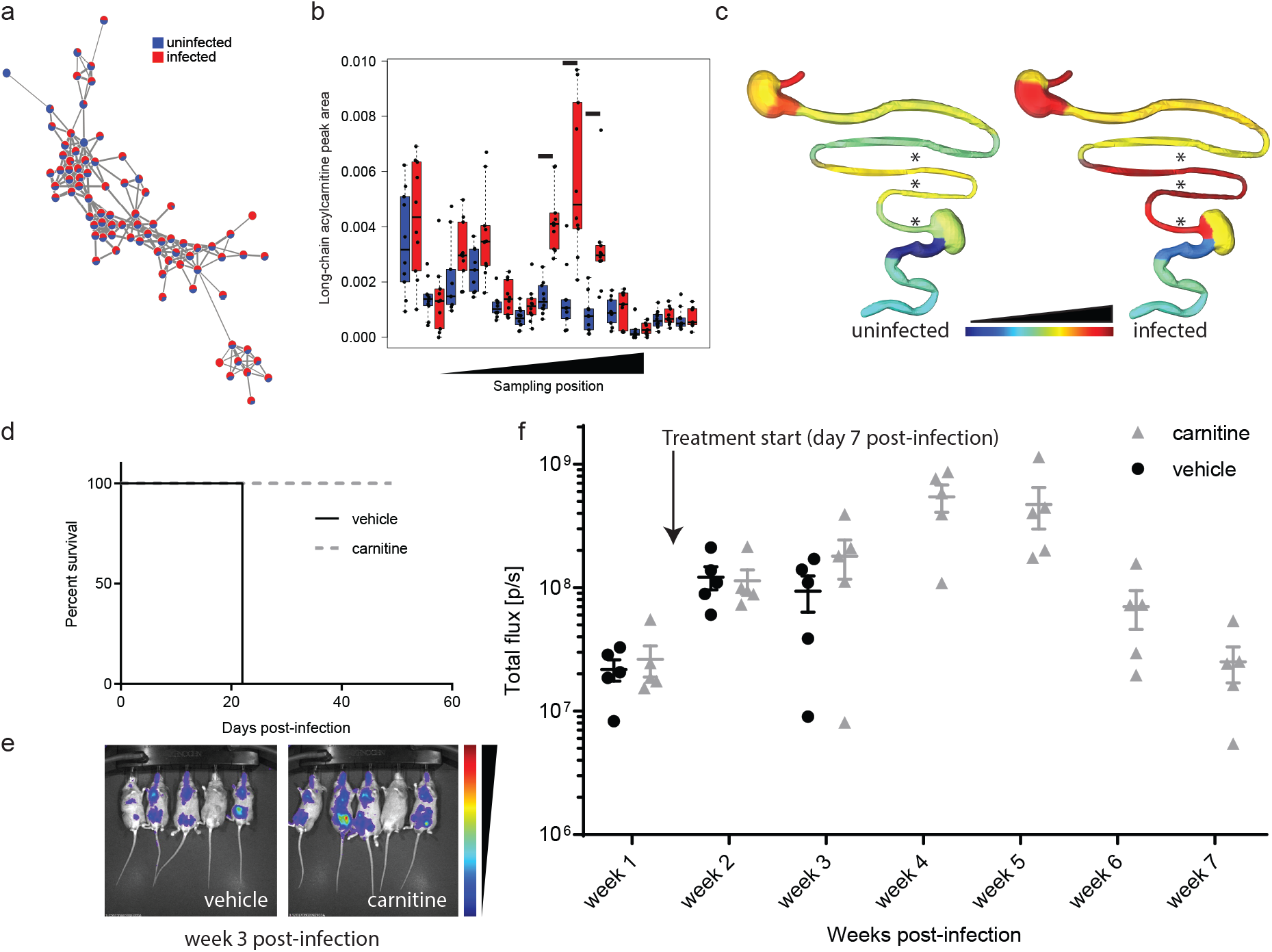
Chemical cartography reveals a causal role for carnitine metabolism in acute CD tolerance. (a) Acylcarnitine molecular network, showing relative abundance of each detected acylcarnitine chemical family member (all tissue sites and timepoints combined; MS2 spectral count relative abundance). Each node represents one metabolite feature. Connected nodes are structurally similar (MS2 cosine score > 0.7), with edge thickness proportional to the cosine score. (b) Infection-induced increases in long-chain acylcarnitines in the distal small intestine 12 days post-infection (black lines indicate FDR-corrected Mann-Whitney p<0.05; p=0.000141, p=0.000563, p=0.000563 for positions 7, 8, 9, comparing infected and uninfected samples for the same positions) (c) Spatial distribution of median long-chain acylcarnitine peak area, 12 days post-infection (common logarithmic scale). Stars indicate sites of statistical significance, as displayed in panel b. (d) Carnitine treatment prevents acute-stage mortality. Male C3H/HeJ mice (n=5 per group) were infected with 50,000 luciferase-expressing *T. cruzi* strain CL Brener. Seven days post-infection, mice were switched to carnitine-supplemented drinking water (carnitine group; 1.3%; equivalent to 100 mg/kg/day based on water consumption) or continued on normal drinking water (vehicle group). All vehicle-treated mice reached humane endpoints at 22 days post-infection, whereas carnitine-treated mice survived the acute infection stage (p=0.0027, Mantel-Cox test). (e) and (f) Comparable parasite burden was observed between carnitine-treated and vehicle groups, indicating that carnitine’s pro-survival effects represent disease tolerance rather than antiparasitic efficacy. (e) Representative bioluminescent imaging, week 3 post-infection (common scale). (f) Overall whole-body luminescence, weeks 1-7 post-infection. Mean and standard error of mean are displayed.

To determine whether we could translate these findings towards novel CD therapeutics and whether these acylcarnitine alterations play a causal role in disease progression, we assessed whether carnitine supplementation could alter acute CD outcome. Mice were infected with either an intermediate dose (5,000 trypomastigotes) or a high dose (50,000 trypomastigotes) of luciferase-expressing CL Brener parasites ^6^. At 7 days post-infection, animals were distributed into two groups of comparable parasite burden and one group switched from normal drinking water to drinking water supplemented with L-carnitine (equivalent to 100 mg/kg/day based on water consumption). Carnitine treatment completely abrogated acute CD-induced mortality up to 7 weeks post-infection (**Fig. 4d**, p=0.0027 Mantel-Cox test, 50,000 trypomastigote infection; **Fig. S12a**), but without any effect on parasite burden or parasite distribution (**Fig. 4ef**, **S12b**). This lack of antiparasitic activity is consistent with prior *in vitro* activity data showing no impact of carnitine on parasite burden ^21^. Overall, these results indicate that acylcarnitine modulation by carnitine supplementation can induce disease tolerance in CD. These results have important implications for our understanding of the factors that contribute to the progression from asymptomatic to symptomatic CD, and represent a novel avenue for CD drug development, in conjunction with antiparasitics to kill *T. cruzi.*

## Discussion

Disease severity is tied to the balance between resistance and tolerance mechanisms ^22^. Resistance reduces pathogen load, but can cause collateral damage to the host, as indeed has been observed with immune clearance of *T. cruzi*-infected cells ^23^. In contrast, tolerance reduces disease or immune-mediated collateral damage without affecting the pathogen load ^22^. While parasite persistence is required for progression to chronic CD ^24^, only a minority of infected patients progress to symptomatic disease ^4^, and parasite load does not fully predict disease severity (*e.g.* ^*25*^), indicating that disease tolerance mechanisms also regulate CD progression, although these are not well understood. Our novel finding that carnitine modulation determines infection outcome (**Fig. 4**) paves the way for future studies of the role of acylcarnitines in the progression from asymptomatic to symptomatic disease in humans, as well as the development of novel interventional strategies for CD, most likely in combination with antiparasitic agents. Our observations also represent the first time that carnitine metabolism has been directly linked to disease tolerance mechanisms, rather than serving as a readout for altered fatty acid metabolism. Importantly, acylcarnitines, as with many other infection-modulated metabolites in our dataset, showed strong spatial effects that would have been masked by bulk tissue analysis, demonstrating the strength of this spatially-resolved approach (**Fig. 2, Fig. 4b**, **Fig. S11**). Strikingly, sites of largest statistically significant overall metabolic disturbance in the chronic stage were the oesophagus and large intestine (**Fig. 1**), providing a mechanism whereby persistent metabolic alterations at these sites drive the striking selective tropism of CD for the large intestine and oesophagus. In contrast, chronic parasite persistence in the cecum was metabolically silent (**Fig. 1**), while cecal microbiome remained strongly and significantly affected by infection (**Fig. 3**). It is tempting to speculate a microbiota-mediated mechanism of reduced antiparasitic immune responses in the cecum, perhaps via cecal microbiota-derived short-chain fatty acid ^26^, or induction of parasite dormancy at this site ^27^, leading to the observed parasite recrudescence in the cecum following incomplete posaconazole or nifurtimox treatment ^8,28^, and this awaits further experimentation. Lastly, the spatially-resolved metabolomic and microbiome methods that we illustrate here with *T. cruzi* can readily be applied to study other pathogens with specific tissue tropism, and we anticipate this approach to have broad applicability. Likewise, initial pathogen tropism is affected by tissue characteristics. Our comprehensive spatial characterization of the microbiome and metabolome of uninfected animals therefore represents a resource that can serve as a hypothesis-generating starting point for studies of pathogen tropism.

## Methods

### *In vivo* experimentation

All vertebrate animal studies were performed in accordance with the USDA Animal Welfare Act and the Guide for the Care and Use of Laboratory Animals of the National Institutes of Health. The protocol was approved by the University of California San Diego Institutional Animal Care and Use Committee (protocol S14187).

For chemical cartography and 16S analysis: 5-week-old male C3H/HeJ mice (The Jackson Laboratory) were infected by intraperitoneal injection of 1,000 red-shifted luciferase-expressing *T. cruzi* strain CL Brener ^6^ culture-derived trypomastigotes in 100 μL DMEM media (infected group) or mock-infected by injection of 100 μL DMEM media only (uninfected group). Prior to animal infection, *T. cruzi* parasites were maintained in coculture with C2C12 mouse myoblasts, in DMEM (Invitrogen) supplemented by 5% iron-supplemented calf serum (HyClone) and 1% penicillin-streptomycin (Invitrogen). Twelve or 89 days post-infection, animals were injected with 150 mg/kg D-luciferin potassium salt (Gold Biotechnology) and euthanized by isoflurane overdose followed by cervical dislocation. Mice were then immediately perfused with 10 mL of 0.3 mg/mL ice-cold D-luciferin in PBS ^6^. GI organs were collected, sectioned as displayed on **Fig. S1a** and each section placed in an individual 96-well-plate well containing 0.3 mg/mL ice-cold D-luciferin in PBS ^6^. The plate was imaged in an *In vivo* Imaging System (IVIS) Lumina LT Series III (Perkin Elmer) and tissues were then immediately snap-frozen in liquid nitrogen, followed by storage at −80 °C. Tissue section luminescence was determined using Living Image 4.5 software, normalized to collected tissue weight, and plotted using GraphPad Prism version 8. Two biological replicate experiments were performed, each including n=5 mice for each timepoint and infection condition (total n=10 per timepoint and infection condition). The same samples were used as source material for 16S and LC-MS analysis (see below), with each tissue site from each individual animal representing a single data point in each analysis.

For carnitine supplementation experiments: mice were infected with either an intermediate dose (5,000 culture-derived trypomastigotes) or a high dose (50,000 culture-derived trypomastigotes) of red-shifted luciferase-expressing CL Brener parasites ^6^. Seven days post-infection, mice were injected with 150 mg/kg D-luciferin potassium salt (Gold Biotechnology) and imaged (IVIS Lumina LT Series III). Animals were allocated to treatment groups to have comparable total body luminescence signal between groups. Mice then received L-carnitine (VWR) in drinking water *ad libitum*, normalized to mouse water consumptions so that animals received ca. 100 mg/kg/day, or regular drinking water (n=5 per group). Bioluminescent imaging was performed weekly. Animals reaching humane endpoints of weight loss >20% were euthanized. Bioluminescence data was analyzed with Living Image 4.5 software and plotted using GraphPad Prism version 8.

### Sample preparation for LC-MS/MS

Samples from both biological replicate experiments were analyzed jointly. Metabolites were extracted from the collected tissue samples using a two-step process as implemented in our prior work ^13^, normalizing to tissue weight. Tissue samples were homogenized in LC-MS grade water (50 mg tissue in 125 μL water) using a 5 mm steel ball in Qiagen TissueLyzer at 25 Hz for 3 min. 10 μL was set aside for DNA extraction and microbiome profile analysis, except for oesophagus where the tissue amount was too small. LC-MS grade methanol spiked with 4 μM sulfachloropyridazine was added to the homogenized sample, to a final concentration of 50% methanol, and the sample was homogenized again at 25 Hz for 3 min. Homogenate was centrifuged for 15 min at 14,980g, 4 °C. The centrifugation supernatant was collected and dried in a Savant SPD111V (ThermoFisher Scientific) speedvac concentrator. The centrifugation pellet was resuspended in 3:1 (by volume) dichloromethane/methanol solvent mixture and further homogenized at 25 Hz for 5 minutes, followed by centrifugation at 14,980g for 2 minutes. This latter centrifugation supernatant was collected and air dried. Both extracts were stored at −80 °C until LC-MS analysis.

### LC-MS/MS

The dried samples were resuspended in 50% methanol (LC-MS grade) spiked with 2μM sulphadimethoxine as internal control, pooling aqueous and organic extracts together. Liquid chromatography was performed using a ThermoScientific Vanquish UHPLC system fitted with 1.7 μm 100 Å Kinetex C8 column (50 × 2.1 mm) (Phenomenex). Data-dependent MS/MS (ddMS^2^) experiments were performed on a Q Exactive Plus (ThermoScientific) high resolution mass spectrometer, under the control of XCalibur and Tune software (ThermoScientific). Ions were generated for MS/MS analysis in both positive and negative ion mode using heated electrospray ionization (HESI) source. Calibration of the instrument was performed using recommended commercial calmix from ThermoScientific. See supplemental information (**Table S4-7**) for detailed instrumental parameters.

### LC-MS/MS data analysis

Raw data was converted to mzXML format using MS Convert software ^29,30^. Processing of the resulting mzXML files was done in MZmine version 2.30 ^31^ (see **Table S8** for parameters). Data was filtered to only retain MS1 scans that were present in at least 6 samples and were associated with MS2 spectra (and therefore could potentially be annotated). Blank removal was performed, with a minimum three-fold difference between blank and samples required in order for a feature to be retained. Total Ion Current (TIC) normalization was performed in Jupyter notebook using R (http://jupyter.org). Principal coordinate analysis (PCoA) was performed on the TIC-normalized MS1 data using the Bray-Curtis-Faith dissimilarity metric in QIIME 1 ^32^, visualized using Emperor ^33^. PERMANOVA calculations were performed on Bray-Curtis-Faith distance matrices using the R package “vegan” ^34^. The 3D GI tract model was built to scale from pictures of GI tract samples collected from our mice, using SketchUp 2017 software. Data was plotted onto this 3D model using ‘ili (https://ili.embl.de/) ^35^. Feature annotation was performed through molecular networking on the Global Natural Products Social Networking (GNPS) platform ^17^ (see **Table S9** for detailed parameters). All annotations are levels 2 or 3 according to the metabolomics standards initiative ^36^. Molecular networks were visualized using Cytoscape ^37,38^. Venn diagrams were generated using: http://bioinformatics.psb.ugent.be/webtools/Venn/. Random forest analysis ^39^ was performed in Jupyter notebook using the randomForest R package and 7501 trees, classifying based on infected vs uninfected status. All code can be accessed at https://github.com/mccall-lab-OU/GI-tract-paper.

### 16S method and data analysis

DNA was extracted from homogenized tissue samples using the DNeasy PowerSoil kit (Qiagen) following manufacturers protocols. The V4 hypervariable region of the 16S rRNA gene was amplified using barcoded Illumina-compatible primers 515F and 806R as previously described ^40^. The resulting amplicons were pooled in equimolar proportions and sequenced on an Illumina MiSeq Instrument. Paired end sequencing reads were quality filtered and merged to reconstruct the complete V4 region using AdapterRemovalV2 ^41^. These analysis-ready reads were used to identify operational taxonomic units (OTUs) following the UNOISE pipeline implemented in Usearch v10 ^42^. Taxonomy was assigned to the representative OTUs using the EzTaxon database ^43^. The resulting OTU table was rarefied to a depth of 5,000 reads per individual, and all downstream statistical analyses were performed using this rarefied OTU table. Alpha- (observed species) and beta-diversity (unweighted UniFrac) analyses were performed using scripts implemented in QIIME 1 ^32^. Kruskal-Wallis tests with FDR correction were used for comparison of genus-level taxonomic summaries to infection status and disease stage.

### Additional statistical information

All statistical tests are paired. Non-parametric tests were used where possible (Mann-Whitney U-test), which makes no assumptions as to data normality. No additional tests of normality were performed. For acylcarnitine data analysis and Figure 2 panels, where Mann-Whitney U-tests were performed for each sampling site, FDR correction was performed to adjust for multiple comparison, as specified in the text and in figure legends. Boxplots display first quartile, median and third quartile, with whiskers no more than 1.5 times interquartile range.

## Data Availability

Metabolomics data have been deposited on MassIVE, accession numbers MSV000082614 (positive mode, oesophagus), MSV000082615 (negative mode, oesophagus), MSV000082618 (positive mode, stomach), MSV000082619 (negative mode, stomach), MSV000082612 (positive mode, small intestine), MSV000082613 (negative mode, small intestine), MSV000082616 (positive mode, large intestine and cecum), and MSV000082617 (negative mode, large intestine and cecum). 16S data has been deposited in the NIH Short Read Archive, project number PRJNA553060. Molecular networks can be accessed here: https://gnps.ucsd.edu/ProteoSAFe/status.jsp?task=801f2cc53c504fad8e64a08565173309# (positive mode networking from mzXML files; used for annotations and dataset matching); https://gnps.ucsd.edu/ProteoSAFe/status.jsp?task=4592e7dfd96c440f8885fb312d50e124 (positive mode feature-based molecular networking; used to cluster MS1 data into chemical families); https://gnps.ucsd.edu/ProteoSAFe/status.jsp?task=d86b3d7c69c646479ef2cf5e7a432ba8 (negative mode networking from mzXML files; used for annotations and dataset matching); https://gnps.ucsd.edu/ProteoSAFe/status.jsp?task=a6415577c1c24bc3823c9af5d9b5092c (negative mode feature-based molecular networking; used to cluster MS1 data into chemical families). Interactive 3D maps of metabolomic data for representative animals can be accessed here: https://ili.embl.de/?ftp://massive.ucsd.edu/MSV000082614/updates/2019-07-10_ehossain_cee84dec/other/3Dmodelofmouse2_ili.stl;ftp://massive.ucsd.edu/MSV000082614/updates/2019-07-10_ehossain_cee84dec/other/Pos_m10_ili.csv (mouse 10, infected, acute-stage sample collection) https://ili.embl.de/?ftp://massive.ucsd.edu/MSV000082614/updates/2019-07-10_ehossain_cee84dec/other/3Dmodelofmouse2_ili.stl;ftp://massive.ucsd.edu/MSV000082614/updates/2019-07-10_ehossain_cee84dec/other/pos_m18_ili.csv (mouse 18, uninfected, chronic-stage sample collection). All code can be accessed at https://github.com/mccall-lab-OU/GI-tract-paper.

## Acknowledgements

This work was supported by start-up funds from the University of Oklahoma to LIM. Initial tissue collection was supported by a postdoctoral fellowship to LIM from the Canadian Institutes of Health Research, award number 338511 (www.cihr-irsc.gc.ca/). Microbial community analysis was supported in part by a National Institutes of Health grant, award number NIH 2R01-GM089886 to KS. The authors wish to thank Dr. James McKerrow and Dr. Jair Lage Siqueira-Neto (UCSD) for advice during the early stages of this project. Dr. John Kelly, London School of Hygiene & Tropical Medicine and Dr. Bruce Branchini, Connecticut College provided the red-shifted luciferase-expressing *T. cruzi* strain CL Brener used in these experiments.

## Author contributions

LIM designed the project. EH and LIM performed metabolite extractions and LC-MS instrumental analysis. EH, CW, DL, MK, CG and LIM performed LC-MS data analysis. DL built the 3D GI tract model. SLJ and DT performed *in vivo* experimentation, carnitine treatment, and tissue sample collection. SK and CMW performed DNA extractions and 16S library builds. KS performed the16S sequencing and analysis. LIM, EH and KS wrote the paper.

