## Supplemental information for "3D mapping of host-parasite-microbiome interactions reveals metabolic determinants of tissue tropism and disease tolerance in Chagas disease"

Affiliations:
[1]Department of Chemistry and Biochemistry, University of Oklahoma, Norman, Oklahoma, United States of America
[2]Laboratories of Molecular Anthropology and Microbiome Research, University of Oklahoma, Norman, Oklahoma, United States of America
[3]Department of Microbiology and Plant Biology, University of Oklahoma, Norman, Oklahoma, United States of America
[4]Skaggs School of Pharmacy and Pharmaceutical Sciences, University of California San Diego, La Jolla, California, United States of America
[5]College of Chemistry, Beijing Normal University, Beijing, China

**Tables S1 to S9**
**Figures S1 to S12**

**Table S1. Infection-modulated metabolites identified through random forest analysis.**

| **12 days post-infection** | | | | | | | | | |
|---|---|---|---|---|---|---|---|---|---|
| **Organ** | ***m/z*** | **RT (min)** | **Annotation** | **Cosine Score** | **Matched Peaks** | **ppm Error** | **Mass Difference to Library Reference** | **Adduct** | **Impact of infection** |
| **Oesophagus** | 448.341 | 2.901 | C20:4 acylcarnitine (spectral match to palmitoylcarnitine) | 0.83 | 7 | 3.79 | 48 | $[M+H]^+$ | increased |
| | 153.041 | 0.441 | - | - | - | - | - | - | increased |
| | 72.081 | 0.36 | - | | - | - | - | - | increased |
| | 86.096 | 0.435 | - | - | - | - | - | - | increased |
| | 269.247 | 3.158 | - | - | - | - | - | - | decreased |
| | 269.136 | 2.381 | - | - | - | - | - | - | decreased |
| | 570.355 | 2.958 | LPC(22:5) (spectral match to LPC(16:0)) | 0.95 | 12 | 0.70 | 74.03 | $[M+H]^+$ | increased |

| | | | | | | | | | |
|---|---|---|---|---|---|---|---|---|---|
| | 176.066 | 0.295 | - | - | - | - | - | - | increased |
| | 329.268 | 3.015 | glyceryl palmitoleate (spectral match to monopalmitolein) [a] | 0.89 | 57 | 3.64 | 18.01 | $[M+H]^+$ | decreased |
| | 421.317 | 2.83 | - | - | - | - | - | - | increased |
| | 138.052 | 0.282 | - | - | - | - | - | - | increased |
| | 568.339 | 2.899 | LPC(22:6) (spectral match to LPC(16:0)) | 0.95 | 14 | 1.41 | 72.01 | $[M+H]^+$ | increased |
| | 299.258 | 3.042 | - | - | - | - | - | - | decreased |
| | 384.394 | 2.716 | - | - | - | - | - | - | increased |
| | 544.339 | 2.902 | LPC(20:4) (spectral match to LPC(16:0)) | 0.94 | 10 | 1.47 | 48.01 | $[M+H]^+$ | increased |
| **Stomach** | 388.305 | 2.732 | C14:0-OH acylcarnitine (spectral match to lauroylcarnitine) | 0.91 | 5 | 1.8 | 44.02 | $[M+H]^+$ | increased |
| | 595.348 | 2.754 | - | - | - | - | - | - | increased |
| | 209.092 | 0.627 | kynurenine (spectral match) | 0.94 | 7 | 2.87 | 0 | $[M+H]^+$ | increased |

| | | | | | | | | | |
|---|---|---|---|---|---|---|---|---|---|
| | 442.352 | 2.853 | C18:1-OH acylcarnitine (spectral match to lauroylcarnitine) | 0.9 | 6 | 2.71 | 98.07 | $[M+H]^+$ | increased |
| | 233.149 | 0.818 | thr-leu (spectral match) | 0.93 | 8 | 0 | 0 | $[M+H]^+$ | decreased |
| | 332.217 | 1.309 | - | - | - | - | - | - | decreased |
| | 440.336 | 2.823 | - | - | - | - | - | - | increased |
| **Small Intestine** | 510.391 | 3.084 | LPC(O-18:0) (spectral match to LPC(16:0)) | 0.97 | 12 | 2.74 | 14.03 | $[M+H]^+$ | increased |
| | 482.36 | 2.967 | LPC(O-16:0) (spectral match to LPC (O-18:0)) | 0.93 | 10 | 1.04 | 28.03 | $[M+H]^+$ | increased |
| | 508.376 | 2.998 | LPC(O-18:1) (spectral match to 1-(1Z-Hexadecenyl)-sn-glycero-3-phosphocholine) | 0.92 | 9 | 1.38 | 28.03 | $[M+H]^+$ | increased |
| | 564.437 | 3.26 | LPC(O-22:1) (spectral match to LPC(O-18:0)) | 0.77 | 8 | 4.07 | 42.04 | $[M+H]^+$ | increased |
| | 550.386 | 3.08 | LPC(20:1) (spectral match to LPC(16:0)) | 0.96 | 11 | 1.27 | 54.06 | $[M+H]^+$ | decreased |

| | | | | | | | | | |
|---|---|---|---|---|---|---|---|---|---|
| | 462.297 | 2.927 | - | - | - | - | - | - | increased |
| | 538.422 | 3.237 | LPC(O-20:0) (spectral match to LPC(O-18:0)) | 0.96 | 12 | 2.04 | 42.06 | $[M+H]^+$ | increased |
| | 448.341 | 2.901 | C20:4 acylcarnitine | 0.83 | 7 | 3.79 | 48 | $[M+H]^+$ | increased |
| | 595.348 | 2.754 | - | - | - | - | - | - | increased |
| | 238.045 | 0.246 | - | - | - | - | - | - | increased |
| | 232.154 | 1.002 | C4:0 acylcarnitine (spectral match to acetylcarnitine) | 0.92 | 4 | 0 | 28.03 | $[M+H]^+$ | increased |
| | 204.123 | 0.282 | acetylcarnitine | 0.94 | 12 | 2.94 | 0 | $[M+H]^+$ | increased |
| | 438.297 | 2.971 | - | - | - | - | - | - | increased |
| | 440.314 | 2.947 | - | - | - | - | - | - | increased |
| **Cecum** | 373.273 | 2.712 | cholic acid (spectral match) | 0.85 | 13 | 8 | 0 | $[M+H-2H_2O]^+$ | increased |
| | 344.716 | 2.197 | - | - | - | - | - | - | increased |
| | 160.133 | 0.271 | - | - | - | - | - | - | increased |

| | | | | | | | | | |
|---|---|---|---|---|---|---|---|---|---|
| | 407.279 | 2.681 | oxocholic acid (spectral match to cholic acid) | 0.76 | 12 | 1.72 | 2.02 | $[M+H]^+$ | increased |
| | 389.268 | 2.662 | - | - | - | - | - | - | increased |
| | 570.355 | 2.958 | LPC(22:5) (spectral match to LPC(16:0)) | 0.95 | 12 | 0.7 | 74.03 | $[M+H]^+$ | increased |
| | 352.541 | 2.168 | - | - | - | - | - | - | increased |
| | 355.245 | 2.621 | - | - | - | - | - | - | decreased |
| | 480.308 | 2.982 | - | - | - | - | - | - | decreased |
| | 209.019 | 0.252 | - | - | - | - | - | - | increased |
| | 214.059 | 0.257 | - | - | - | - | - | - | increased |
| | 406.233 | 2.525 | - | - | - | - | - | - | increased |
| | 127.05 | 0.711 | - | - | - | - | - | - | increased |
| | 159.076 | 0.255 | - | - | - | - | - | - | increased |

| | | | | | | | | | |
|---|---|---|---|---|---|---|---|---|---|
| **Large Intestine** | 209.092 | 0.627 | kynurenine (spectral match) | 0.94 | 7 | 2.87 | 0 | $[M+H]^+$ | increased |
| | 572.37 | 2.99 | LPC(22:4) (spectral match to LPC(16:0)) | 0.94 | 12 | 1.92 | 76.04 | $[M+H]^+$ | increased |
| | 119.09 | 0.344 | - | - | - | - | - | - | decreased |
| | 188.071 | 1.187 | tryptophan (spectral match) | 0.93 | 5 | 0 | 0 | $[M+H-NH_3]^+$ | decreased |
| | 570.355 | 2.958 | LPC(22:5) (spectral match to LPC(16:0)) | 0.95 | 12 | 0.7 | 74.03 | $[M+H]^+$ | increased |
| | 205.097 | 1.243 | tryptophan (spectral match) | 0.98 | 7 | 5 | 0 | $[M+H]^+$ | decreased |
| | 160.133 | 0.271 | - | - | - | - | - | - | increased |
| | 153.041 | 0.441 | - | - | - | - | - | - | increased |
| | 544.339 | 2.902 | LPC(20:4) (spectral match to LPC(16:0)) | 0.94 | 10 | 1.47 | 48.01 | $[M+H]^+$ | increased |
| **89 days post-infection** | | | | | | | | | |
| **Organ** | ***m/z*** | **RT (min)** | **Annotation** | **Cosine Score** | **Matched Peaks** | **ppm Error** | **Mass Difference to Library Reference** | **Adduct** | **Impact of infection** |

| | | | | | | | | | |
|---|---|---|---|---|---|---|---|---|---|
| **Oesophagus** | 148.06 | 0.365 | - | - | - | - | - | - | increased |
| | 427.266 | 2.567 | - | - | - | - | - | - | decreased |
| | 120.065 | 0.364 | - | - | - | - | - | - | increased |
| | 566.321 | 2.908 | LPC(20:4) (spectral match to LPC(O-18:0)) | 0.88 | 7 | 1.24 | 0 | $[M+Na]^+$ | increased |
| | 570.355 | 2.958 | LPC(22:5) (spectral match to LPC(16:0)) | 0.95 | 12 | 0.7 | 74.03 | $[M+H]^+$ | increased |
| | 544.339 | 2.902 | LPC(20:4) (spectral match to LPC(16:0)) | 0.94 | 10 | 1.47 | 48.01 | $[M+H]^+$ | increased |
| | 404.3 | 2.767 | - | - | - | - | - | - | decreased |
| | 651.205 | 2.355 | - | - | - | - | - | - | increased |
| | 150.058 | 0.375 | methionine (spectral match) | 0.95 | 6 | 0 | 0 | $[M+H]^+$ | increased |
| | 568.339 | 2.899 | LPC(22:6) (spectral match to LPC(16:0)) | 0.95 | 14 | 1.41 | 72.01 | $[M+H]^+$ | increased |
| | 162.112 | 0.258 | carnitine (spectral match) | 0.75 | 4 | 6 | 0 | $[M+H]^+$ | increased |
| **Stomach** | 167.031 | 0.319 | - | - | - | - | - | - | increased |

| | | | | | | | | | |
|---|---|---|---|---|---|---|---|---|---|
| | 134.045 | 0.397 | - | - | - | - | - | - | increased |
| | 383.326 | 2.644 | - | - | - | - | - | - | increased |
| | 263.016 | 0.276 | - | - | - | - | - | - | increased |
| | 395.29 | 2.624 | - | - | - | - | - | - | increased |
| | 180.136 | 2.139 | - | - | - | - | - | - | decreased |
| | 326.207 | 2.173 | - | - | - | - | - | - | increased |
| | 595.348 | 2.754 | - | - | - | - | - | - | increased |
| | 117.074 | 0.354 | - | - | - | - | - | - | increased |
| | 379.295 | 2.625 | - | - | - | - | - | - | increased |
| **Small Intestine** | 387.289 | 2.843 | - | - | - | - | - | - | increased |
| | 357.278 | 2.877 | chenodeoxycholic acid (spectral match) [b] | 0.91 | 124 | 0 | 0 | $[M+H-2H_20]^+$ | increased |
| | 817.581 | 2.725 | cholic acid | 0.83 | 12 | 2 | 0 | $[2M+H]^+$ | increased |

| | | | | | | | | | |
|---|---|---|---|---|---|---|---|---|---|
| **Cecum** | 324.289 | 3.089 | - | - | - | - | - | - | increased |
| | 276.163 | 2.273 | - | - | - | - | - | - | decreased |
| | 280.263 | 3.172 | - | - | - | - | - | - | increased |
| | 80.948 | 0.255 | - | - | - | - | - | - | decreased |
| | 538.519 | 4.068 | Cer(d34:1) (spectral match to Cer(d18:1/20:1)) | 0.86 | 7 | 0.74 | 53.04 | $[M+H]^+$ | increased |
| | 520.508 | 4.063 | Cer(d34:1) (spectral match to Cer(d18:1/16:1)) | 0.79 | 6 | 1.55 | 14.99 | $[M+H-H_2O]^+$ | increased |
| | 115.037 | 0.29 | - | - | - | - | - | - | decreased |
| | 572.37 | 2.99 | LPC(22:4) (spectral match to LPC(16:0)) | 0.94 | 12 | 1.92 | 76.04 | $[M+H]^+$ | increased |
| | 138.055 | 2.162 | - | - | - | - | - | - | increased |
| | 282.279 | 3.253 | Cer(C24:1) (spectral match) | 0.81 | 7 | 3 | 0 | $[M+H-C_{24}H_{44}O-H_2O]^+$ | increased |

| | | | | | | | | | |
|---|---|---|---|---|---|---|---|---|---|
| **Large Intestine** | 209.092 | 0.627 | kynurenine (spectral match) | 0.94 | 7 | 2.87 | 0 | $[M+H]^+$ | increased |
| | 279.231 | 2.974 | 9(10)-EpOME (spectral match) [c] | 0.92 | 75 | 2 | 0 | $[M+H-H_2O]^+$ | decreased |
| | 618.304 | 2.559 | - | - | - | - | - | - | decreased |
| | 305.247 | 3.242 | - | - | - | - | - | - | increased |
| | 204.105 | 2.377 | - | - | - | - | - | - | decreased |
| | 282.279 | 3.253 | - | - | - | - | - | - | increased |
| | 295.226 | 2.849 | - | - | - | - | - | | decreased |
| | 293.247 | 3.151 | 9Z,11E,13E-octadecatrienoic acid methyl ester (spectral match) [d] | 0.93 | 57 | 1 | 0 | $[M+H]^+$ | decreased |
| | 280.263 | 3.172 | - | - | - | - | - | - | increased |
| | 572.37 | 2.99 | LPC(22:4) (spectral match to LPC(16:0)) | 0.94 | 12 | 1.92 | 76.04 | $[M+H]^+$ | increased |

[a] Match obtained in the absence of 50 Da window filtering. Data can be accessed here: https://gnps.ucsd.edu/ProteoSAFe/status.jsp?task=e46a139901094b63af54cb0e8371d164

[b] Match obtained in the absence of 50 Da window filtering. Data can be accessed here:

https://gnps.ucsd.edu/ProteoSAFe/status.jsp?task=9480227e00ce42b385ab5a509aa57f81

[c] Match obtained in the absence of 50 Da window filtering. Data can be accessed here: https://gnps.ucsd.edu/ProteoSAFe/status.jsp?task=b3dd83cc7874499f834a30c5c475d90d

[d] Match obtained in the absence of 50 Da window filtering. Data can be accessed here: https://gnps.ucsd.edu/ProteoSAFe/status.jsp?task=68805496317a45d19d8227d9ce751eb5

**Table S2. Representative microbially-modified, microbially-derived and microbially-influenced molecules detected in this dataset.**

| *m/z* | RT (min) | GNPS library annotation | Adduct | Cosine score | Matched peaks | Mass difference | ppm error |
|---|---|---|---|---|---|---|---|
| 188.071 | 2.18 | indole-3-lactic acid | $[M+H-H_2O]^+$ | 0.86 | 6 | 0 | 1.06 |
| 212.002 | 2.28 | indoxyl sulfate | $[M-H]^-$ | 0.99 | 4 | 0 | 0.94 |
| 785.589 | 2.89 | deoxycholic acid | $[2M+H]^+$ | 0.86 | 16 | 0 | 5.22 |
| 205.097 | 1.24 | tryptophan | $[M+H]^+$ | 0.98 | 7 | 0 | 3.41 |
| 165.055 | 0.40 | tyrosine | $[M+H-NH_3]^+$ | 0.94 | 5 | 0 | 1.21 |
| 527.158 | 0.27 | maltotriose | $[M+Na]^+$ | 0.88 | 7 | 0 | 1.52 |

**Table S3. Detected acylcarnitines.**

| **Short-chain acylcarnitines** | | | |
|---|---|---|---|
| ***m/z*** | **RT (min)** | **Putative annotation** | **ppm error** |
| 204.123 | 0.282 | C2:0 acylcarnitine (acetylcarnitine) | 2.94 |
| 218.138 | 0.466 | C3:0 acylcarnitine (propionylcarnitine) | 5.50 |
| 232.154 | 0.349 | C4:0 acylcarnitine (butyrylcarnitine) | 0 |
| 232.154 | 1.002 | C4:0 acylcarnitine (butyrylcarnitine) | 0 |
| 248.149 | 0.404 | C4:0-OH acylcarnitine | 3.22 |
| 248.149 | 0.267 | C4:0-OH acylcarnitine | 3.22 |
| **Mid-chain acylcarnitines** | | | |
| ***m/z*** | **RT (min)** | **Putative annotation** | **ppm error** |
| 246.17 | 2.205 | C5:0 acylcarnitine (valerylcarnitine) | 2.03 |
| 260.185 | 2.304 | C6:0 acylcarnitine | 4.61 |
| 260.186 | 2.383 | C6:0 acylcarnitine | 0.77 |
| 288.216 | 2.514 | C8:0 acylcarnitine | 4.20 |
| 314.233 | 2.599 | C10:1 acylcarnitine | 0.32 |
| 316.247 | 2.645 | C10:0 acylcarnitine | 5.69 |
| 332.243 | 2.546 | C10:0-OH acylcarnitine | 2.11 |
| **Long-chain acylcarnitines** | | | |
| ***m/z*** | **RT (min)** | **Putative annotation** | **ppm error** |
| 342.263 | 2.865 | C12:1 acylcarnitine | 4.09 |
| 344.279 | 2.752 | C12:0 acylcarnitine (lauroylcarnitine) [a] | 3.20 |
| 360.273 | 2.655 | C12:0-OH acylcarnitine | 2.78 |

| | | | |
|---|---|---|---|
| 368.279 | 2.808 | C14:2 acylcarnitine | 2.99 |
| 370.294 | 2.789 | C14:1 acylcarnitine | 4.59 |
| 370.295 | 2.982 | C14:1 acylcarnitine | 1.89 |
| 372.237 | 2.399 | C12:2-DC acylcarnitine | 4.30 |
| 372.310 | 2.998 | C14:0 acylcarnitine | 3.76 |
| 372.310 | 2.835 | C14:0 acylcarnitine | 3.76 |
| 388.305 | 2.732 | C14:0-OH acylcarnitine | 1.80 |
| 396.310 | 2.829 | C16:2 acylcarnitine | 3.53 |
| 400.269 | 2.605 | C14:2-DC acylcarnitine | 2.25 |
| 400.341 | 2.929 | C16:0 acylcarnitine (palmitoylcarnitine) | 4.25 |
| 402.284 | 2.636 | C14:1-DC acylcarnitine | 3.98 |
| 412.305 | 2.72 | C16:2-OH acylcarnitine | 3.15 |
| 414.32 | 2.773 | C16:1-OH acylcarnitine | 4.59 |
| 414.321 | 2.841 | C16:1-OH acylcarnitine | 2.17 |
| 422.326 | 2.848 | C18:3 acylcarnitine | 2.37 |
| 424.341 | 2.896 | C18:2 acylcarnitine | 4.01 |
| 426.357 | 2.953 | C18:1 acylcarnitine (oleoylcarnitine) | 3.05 |
| 428.3 | 2.67 | C16:2-DC acylcarnitine | 2.80 |
| 428.373 | 3.018 | C18:0 acylcarnitine | 2.33 |
| 430.316 | 2.698 | C16:1-DC acylcarnitine | 2.09 |
| 440.336 | 2.823 | C18:2-OH acylcarnitine | 3.63 |
| 442.352 | 2.853 | C18:1-OH acylcarnitine | 2.71 |
| 444.331 | 2.773 | C17:1-DC acylcarnitine | 3.38 |

| | | | |
|---|---|---|---|
| 444.367 | 2.911 | C18:0-OH acylcarnitine | 2.25 |
| 448.341 | 2.901 | C20:4 acylcarnitine | 3.79 |
| 452.373 | 2.977 | C20:2 acylcarnitine | 2.21 |
| 454.316 | 2.706 | C18:3-DC acylcarnitine | 1.98 |
| 454.388 | 3.037 | C20:1 acylcarnitine | 3.52 |
| 456.331 | 2.725 | C18:2-DC acylcarnitine | 3.29 |
| 456.404 | 3.099 | C20:0 acylcarnitine | 2.84 |
| 458.348 | 2.726 | C18:1-DC acylcarnitine | 0.44 |
| 458.348 | 2.76 | C18:1-DC acylcarnitine | 0.44 |
| 512.466 | 3.52 | C24:0 acylcarnitine | 3.71 |

[a] GNPS library match (see **Fig. S11a**).

**Table S4. Liquid Chromatography Gradient.**

| Time (min) | Flow (mL/min) | %B | Curve |
|---|---|---|---|
| | Run | | |
| 0.00 | 0.500 | 2 | 5 |
| 1.00 | 0.500 | 2 | 5 |
| 2.50 | 0.500 | 98 | 5 |
| 4.50 | 0.500 | 98 | 5 |
| 5.50 | 0.500 | 2 | 5 |
| 7.50 | 0.500 | 2 | 5 |
| 7.50 | Stop Run | | |

A = Water + 0.1% Formic Acid
B = Acetonitrile + 0.1% Formic Acid

**Table S5. Heated electrospray ionization (HESI) source parameters (positive and negative mode).**

| | |
|---|---|
| Sheath Gas Flow Rate (L/min) | 35 |
| Aux Gas Flow Rate (L/min) | 10 |
| Sweep Gas Flow Rate (L/min) | 0 |
| Spray Voltage (kV) | 3.80 (+) / 3.0 (-) |
| Capillary Temperature (ºC) | 320 |
| S-lens RF Level | 50.0 |
| Aux Gas Heater Temperature (ºC) | 350.0 |

**Table S6. Instrumental method.**

| **Properties of the method** | |
|---|---|
| Use Lock Masses | Off |
| Chromatogram Peak Width | 6 seconds |
| Method Duration | 7.50 minutes |
| **Properties of Divert Valve A** | |
| Used | True |
| Start in 1-2 | True |
| Switch Count | 1 |
| At | 0.20 minute |
| Switch to | 1-6 |
| **Properties of Full MS / dd-MS$^2$** | |
| *General* | |
| Runtime | 0 to 7.50 minutes |
| Default Charge State | 1 |
| Inclusion | --- |
| Exclusion | On |
| Tag | --- |
| *Full MS* | |
| Resolution | 70,000 |
| AGC Target | 1E6 |
| Maximum IT | 246 milliseconds |
| Scan Range | 70 to 1050 m/z |
| *dd-MS$^2$* | |
| Resolution | 17,500 |
| AGC Target | 2E5 |
| Maximum IT | 54 milliseconds |
| Loop Count | 5 |
| Top N | 5 |
| Isolation Window | 1.0 *m/z* |
| Fixed First Mass | --- |
| (N)CE/ Stepped | NCE: 20, 40, 60 |
| *dd Settings* | |
| Min. AGC Target | 8.00E3 |
| Intensity Threshold | 1.5E5 |
| Apex Trigger | --- |
| Charge Exclusion | --- |
| Peptide Match | preferred |
| Exclude Isotope | On |
| Dynamic Exclusion | 10.0 seconds |

**Table S7. Exclusion lists.**

| *m/z* | **Polarity** |
|---|---|
| 371.101 | Positive / Negative |
| 235.206 | Positive / Negative |
| 311.084 | Positive / Negative |
| 314.131 | Positive / Negative |
| 285.0135 | Positive / Negative |
| 144.9822 | Positive / Negative |

**Table S8. mzMine (version 2.30) parameters.**

| | | **Polarity** | |
|---|---|---|---|
| | | Positive | Negative |
| **$MS^1$** | Retention Time (min) | 0.0 – 7.51 | 0.0 – 7.51 |
| | Noise Level | 2.0E6 | 2.0E5 |
| **$MS^2$** | Retention Time (min) | 0.0 – 7.51 | 0.0 – 7.51 |
| | Noise Level | 1.0E3 | 1.0E3 |
| **Chromatogram Builder** | Mass List | masses | masses |
| | Minimum Time Span (min) | 0.01 | 0.01 |
| | Minimum Height | 6.0E6 | 5.0E5 |
| | *m/z* Tolerance (ppm) | 10.0 | 10.0 |
| **Chromatogram Deconvolution** | Algorithm | Baseline Cut-off | Baseline Cut-off |
| | *m/z* Range for $MS^2$ Scan Pairing (Da) | 0.01 | 0.01 |
| | RT Range for $MS^2$ Scan Pairing (min) | 0.2 | 0.2 |
| | Minimum Peak Height | 6.0E6 | 5.0E5 |
| | Peak Duration Range (min) | 0.01 – 3.50 | 0.01 – 3.50 |
| | Baseline Level | 2.0E6 | 2.0E5 |
| **Deisotoping** | m/z Tolerance (ppm) | 10.0 | 10.0 |
| | Retention Time Tolerance (min) | 0.5 | 0.05 |
| | Monotonic Shape | Checked | Checked |
| | Maximum Charge | 3 | 3 |
| | Representative Isotope | Lowest m/z | Lowest m/z |
| **Alignment** | *m/z* Tolerance (ppm) | 10.0 | 10.0 |
| | Weight for *m/z* | 1 | 1 |
| | Weight for Retention Time | 1 | 1 |
| | Retention Time Tolerance (min) | 0.5 | 0.5 |
| **Row Filtering** | Minimum Peaks in a Row | 6 | 6 |
| | Retention Time (min) | 0.20 – 6.45 | 0.20 – 6.45 |
| | Keeps Only Peaks with $MS^2$ Scans | Checked | Checked |
| | Reset Peak No. ID | Checked | Checked |

**Table S9. GNPS parameters.**

| | |
|---|---|
| Pairs minimum cosine | 0.7 |
| Analog search | yes |
| Precursor ion mass tolerance (Da) | 0.02 |
| Fragment ion mass tolerance (Da) | 0.02 |
| Minimum matched peaks | 4 |
| Topk | 10 |
| Minimum cluster size [a] | 4 |
| Maximum component size | 100 |
| Minimum peak intensity | 0 |
| Filter standard deviation peak intensity | 0 |
| Run MS cluster [a] | on |
| Filter precursor window | yes |
| Filter library | yes |
| 50 Da window filter | yes |
| Score threshold (library search) | 0.7 |
| Minimum matched peaks (library search) | 4 |
| Maximal mass shift | 100 |
| Normalization per file [b] | row sum normalization |
| Aggregation Method For peak abundances per group | sum |
| Top results per query [b] | 1 |

[a] Only for standard molecular networking workflow

[b] Only for feature-based molecular networking workflow

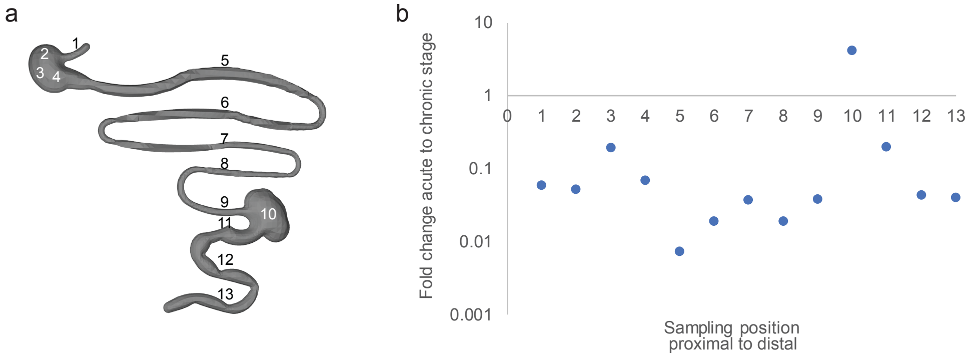

**Fig. S1. Acute-to-chronic stage changes in parasite burden.** (a) Sampling sites. (b) Fold change in median luminescent signal between acute and chronic stages. Parasite burden decreased at all sampling sites except the cecum.

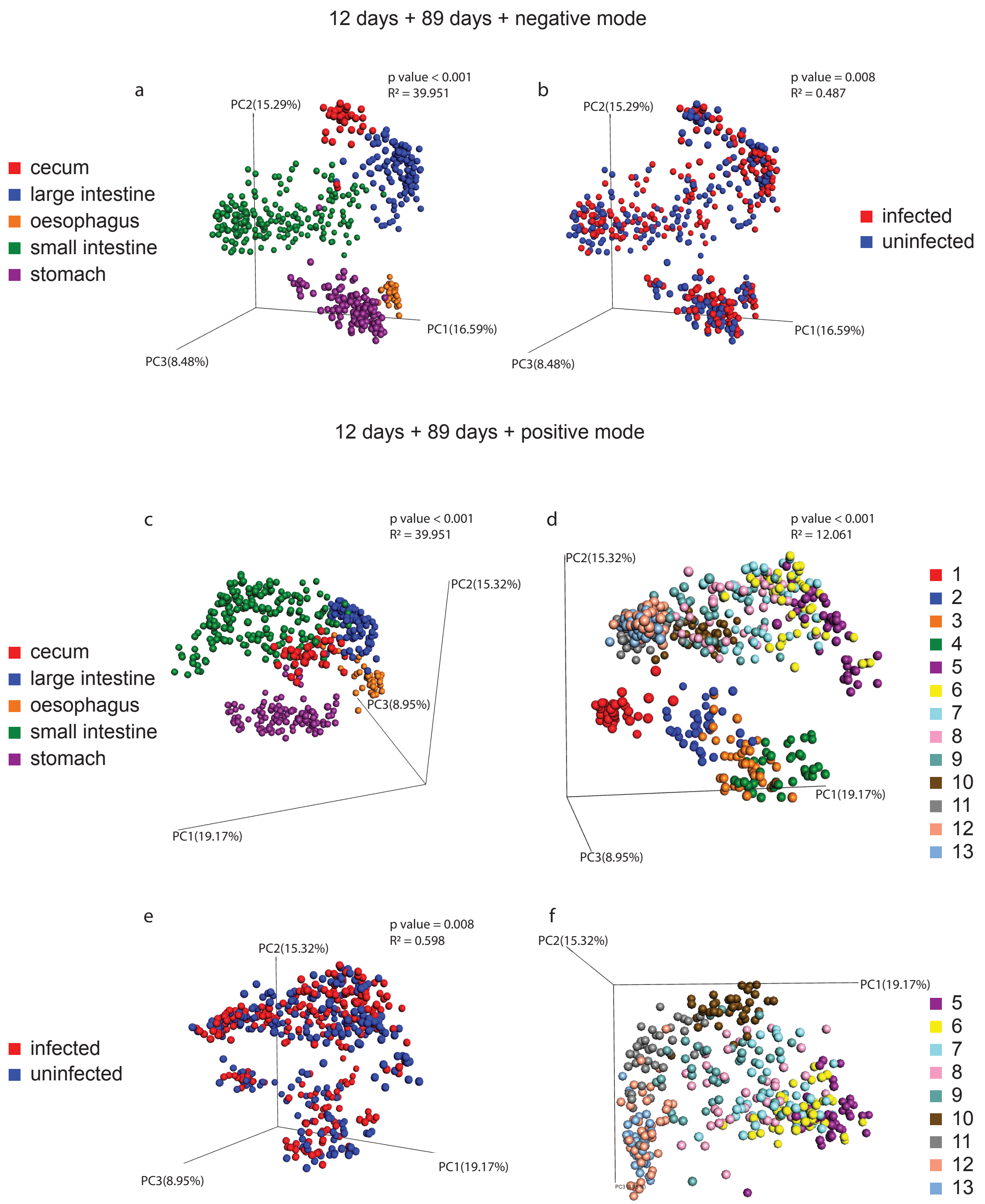

**Fig. S2. Principal coordinate analysis (Bray-Curtis-Faith distance metric) reveals large-scale differences in overall small molecule profile between organs and sampling sites, with minor but significant infection-associated impact, when all timepoints are analyzed jointly.**

(a) and (b), negative mode analysis. (c-f), positive mode analysis. (d) and (f) are views of the same data, with sampling positions 1-4 masked in panel (f) for viewing clarity. (a) and (c) Impact of organ on chemical profile. (d) Impact of sampling site on chemical profile. Refer to **Fig. S1** for sampling position numbers. (b) and (e) Impact of infection status. PERMANOVA $R^2$ and p value are displayed on the plots.

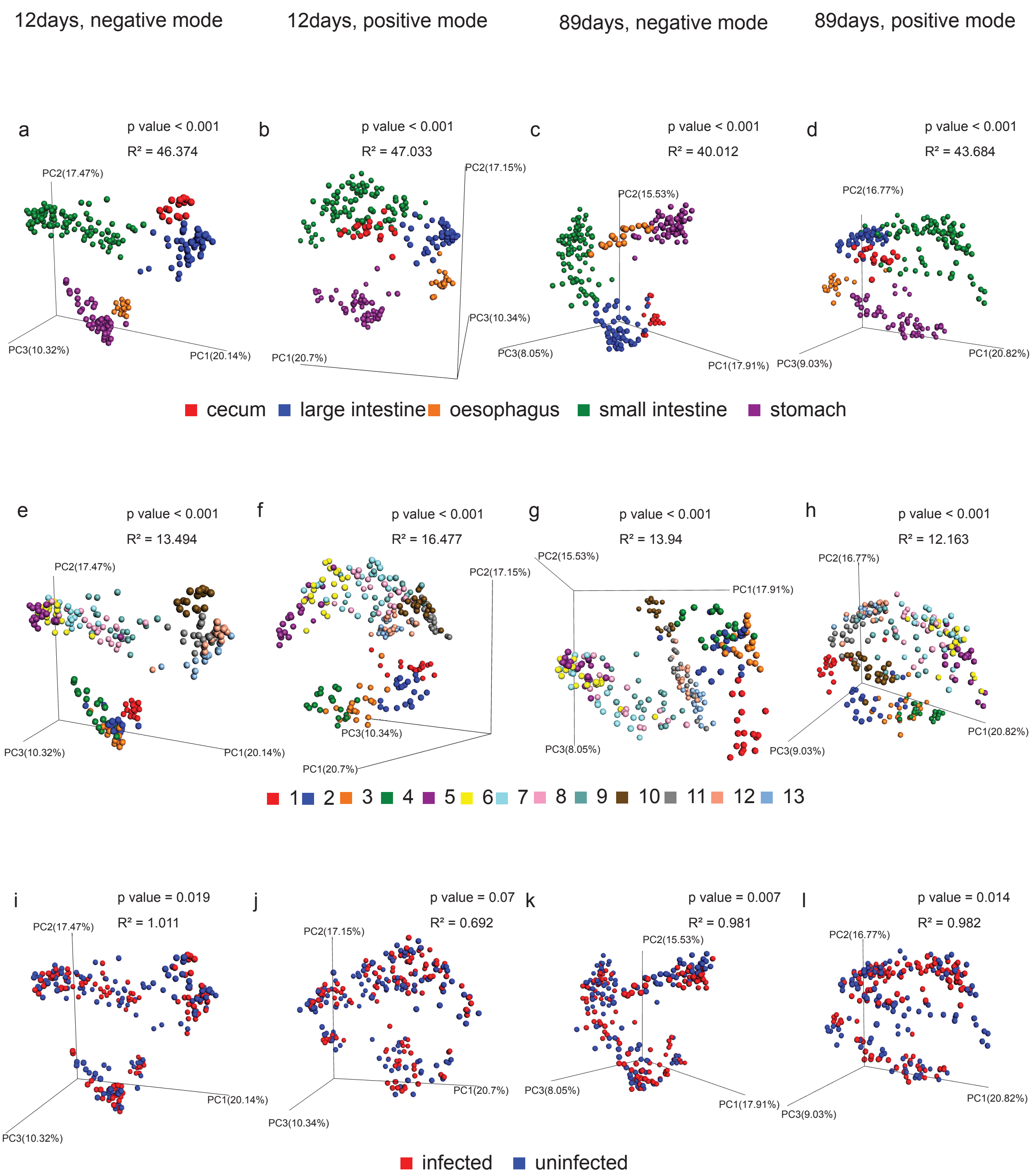

**Fig. S3. Differences in overall chemical composition by organ and by sampling site are maintained when each timepoint is analyzed individually.** Principal coordinate analysis was performed using the Bray-Curtis-Faith distance metric on detected metabolite features. (a), (e), (i), samples collected 12 days post-infection and analyzed in negative mode. (b), (f), (j), samples collected 12 days post-infection and analyzed in positive mode. (c), (g), (k), samples collected 89 days post-infection and analyzed in negative mode. (d), (h), (l), samples collected 89 days post-infection and analyzed in positive mode. (a-d) Impact of organ on chemical profile. (e-h) Impact of sampling site on overall chemical profile. Refer to **Fig. S1** for sampling position numbers. (i-l) Impact of infection status on overall chemical profile. PERMANOVA $R^2$ and p value are displayed on the plots.

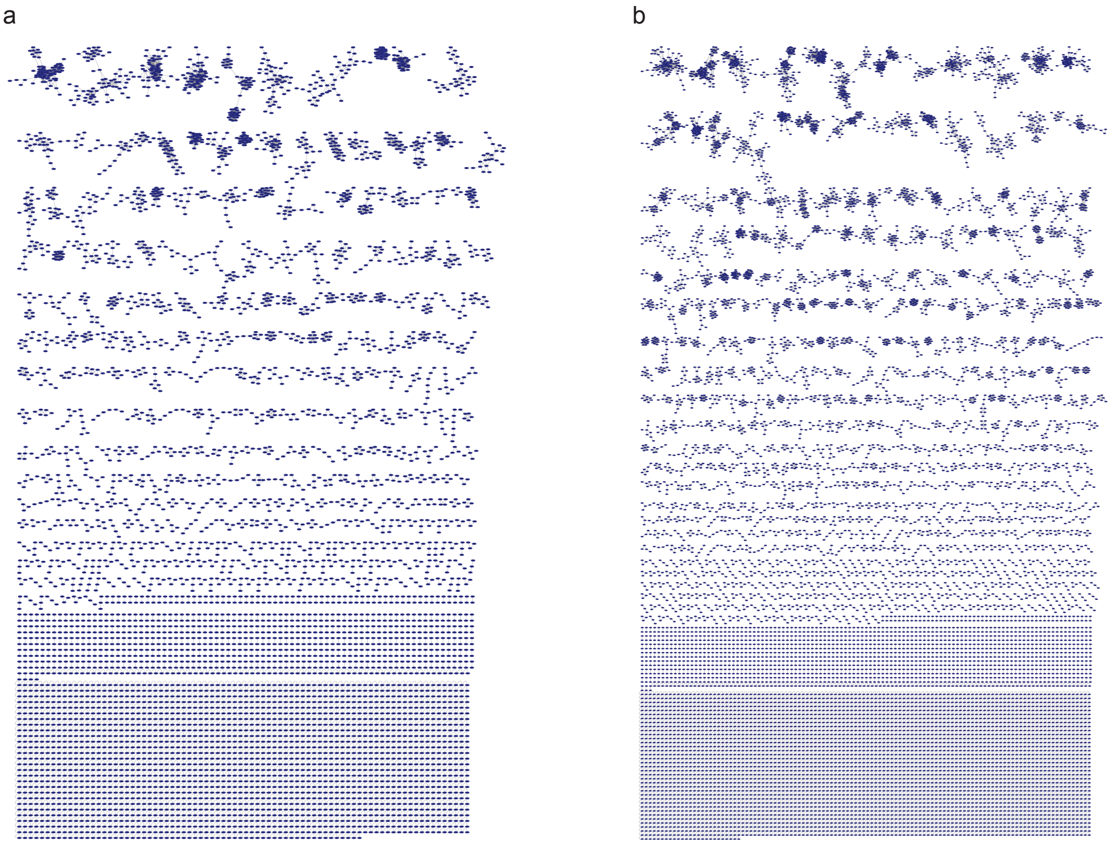

**Fig. S4. Molecular networks.** Networks of chemical families, from feature-based molecular networking. (a) Positive mode. (b) Negative mode.

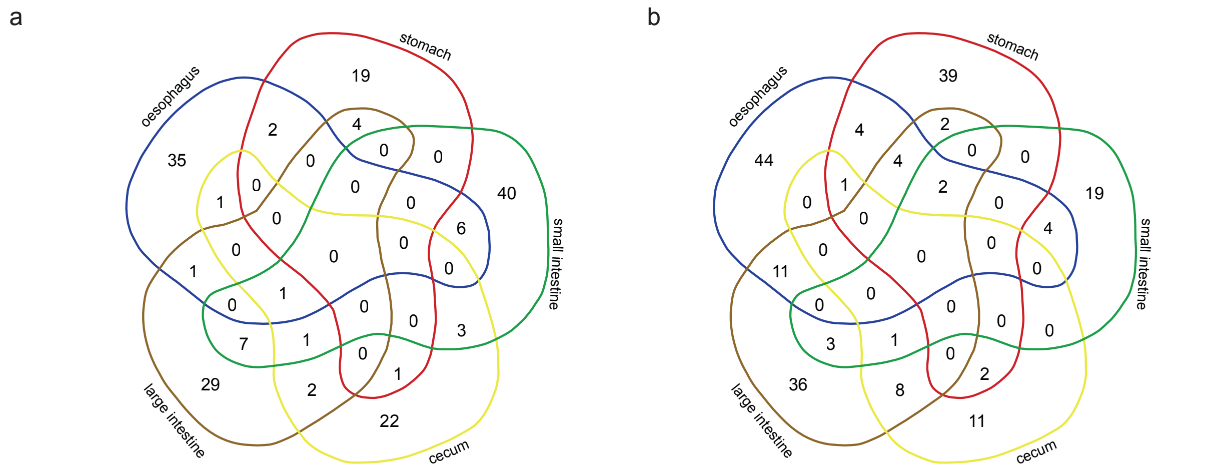

**Fig. S5. Limited overlap in chemical families differentially-modulated by infection, across organs.** Metabolite features were combined into chemical families (as identified by feature-based molecular networking, see **Fig. S4**). Chemical families with fold change in abundance >2 and Mann-Whitney p-value<0.05 were compared between sampling sites. (a) Acute stage. (b) Chronic stage.

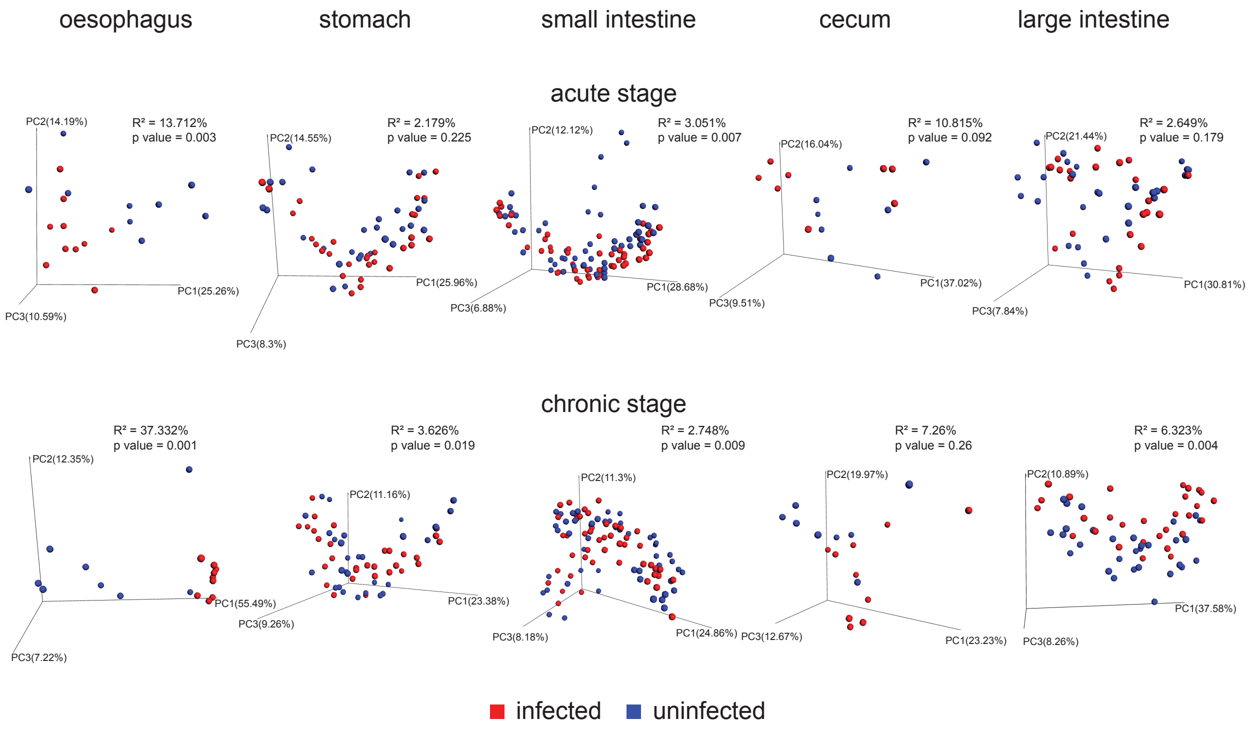

**Fig. S6. Principal coordinate analysis of negative mode data confirms that the largest statistically significant metabolic perturbation in the chronic stage is at the sites of CD pathogenesis, the oesophagus and the large intestine.** Top, acute stage. Bottom, chronic stage.

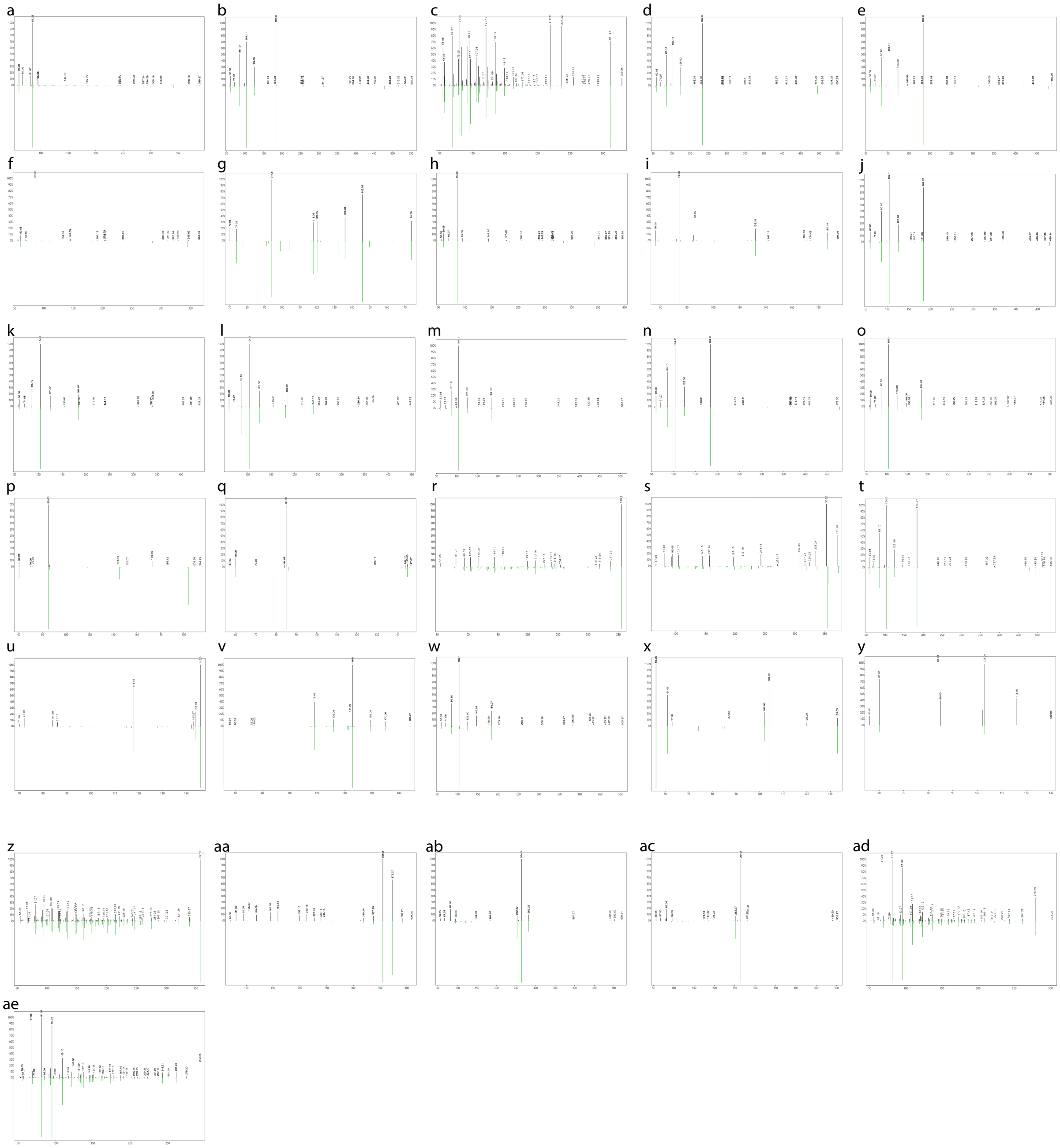

**Fig. S7. Annotation support for infection-modulated molecules identified by random forest analysis.** All annotations are level 2/3 according to the metabolomics standards initiative 36. (a) m/z 488.341 RT 2.90 min (top, black) match to palmitoylcarnitine library reference (bottom, green). (b) m/z 570.355 RT 2.96 min (top, black) match to LPC(16:0) library reference (bottom, green). (c) m/z 329.268 RT 3.02 min (top, black) match to monopalmitolein library reference (bottom, green). (d) m/z 568.339 RT 2.90 min (top, black) match to LPC(16:0) library reference (bottom, green). (e) m/z 544.339 RT 2.90 min (top, black) match to LPC(16:0) library reference (bottom, green). (f) m/z 388.305 RT 2.73 min (top, black) match to lauryolcarnitine library reference (bottom, green). (g) m/z 209.092 RT 0.63 min (top, black) match to L-kynurenine library reference (bottom, green). (h) m/z 442.352 RT 2.85 min (top, black) match to lauryolcarnitine library reference (bottom, green). (i) m/z 233.149 RT 0.82 min (top, black) match to thr-leu library reference (bottom, green). (j) m/z 510.391 RT 3.08 min (top, black) match to LPC(16:0) library reference (bottom, green). (k) m/z 482.36 RT 2.97 min (top, black) match to Lyso PAF C-18 (LPC(O-18:0)) library reference (bottom, green). (l) m/z 508.376 RT 3.00 min (top, black) match to 1-(1Z-Hexadecenyl)-sn-glycero-3-phosphocholine library reference (bottom, green). (m) m/z 564.437 RT 3.26 min (top, black) match to LPC(O-18:0) library reference (bottom, green). (n) m/z 550.386 RT 3.08 min (top, black) match to LPC(16:0) library reference (bottom, green). (o) m/z 538.422 RT 3.24 min (top, black) match to LPC(16:0) library reference (bottom, green). (p) m/z 232.154 RT 1.00 min (top, black) match to acetyl-DL-carnitine library reference (bottom, green). (q) m/z 204.123 RT 0.28 min (top, black) match to acetyl-DL-carnitine library reference (bottom, green). (r) m/z 373.273 RT 2.71 min (top, black) match to cholic acid library reference (bottom, green). (s) m/z 407.279 RT 2.68 min (top, black) match to cholic acid library reference (bottom, green). (t) m/z 572.37 RT 2.99 min (top, black) match to LPC(16:0) library reference (bottom, green). (u) m/z 188.071 RT 1.19 min (top, black) match to tryptophan library reference (bottom, green). (v) m/z 205.097 RT 1.24 min (top, black) match to tryptophan library reference (bottom, green). (w) m/z 566.321 RT 2.91 min (top, black) match to Lyso PAF C-18 (LPC(O-18:0)) library reference (bottom, green). (x) m/z 150.058 RT 0.38 min (top, black) match to 2-amino-4-methylsulfanylbutanoic acid (methionine) library reference (bottom, green). (y) m/z 162.112 RT 0.26 min (top, black) match to L-carnitine library reference (bottom, green). (z) m/z 357.278 RT 2.88 min (top, black) match to 3 beta -hydroxy-5-cholenoic acid library reference (bottom, green). (aa) m/z 817.518 RT 2.73 min (top, black) match to cholic acid library reference (bottom, green). (ab) m/z 538.519 RT 4.07 min (top, black) match to Cer(d18:1/20:1) library reference (bottom, green). (ac) m/z 520.508 RT 4.06 min (top, black) match to Cer(d18:1/16:1) library reference (bottom, green). (ad) m/z 279.231 RT 2.97 min (top, black) match to 9(10)-EpOME library reference (bottom, green). (ae) m/z 293.247 RT 3.15 min (top, black) match to 9Z,11E,13E-octadecatrienoic acid methyl ester library reference (bottom, green).

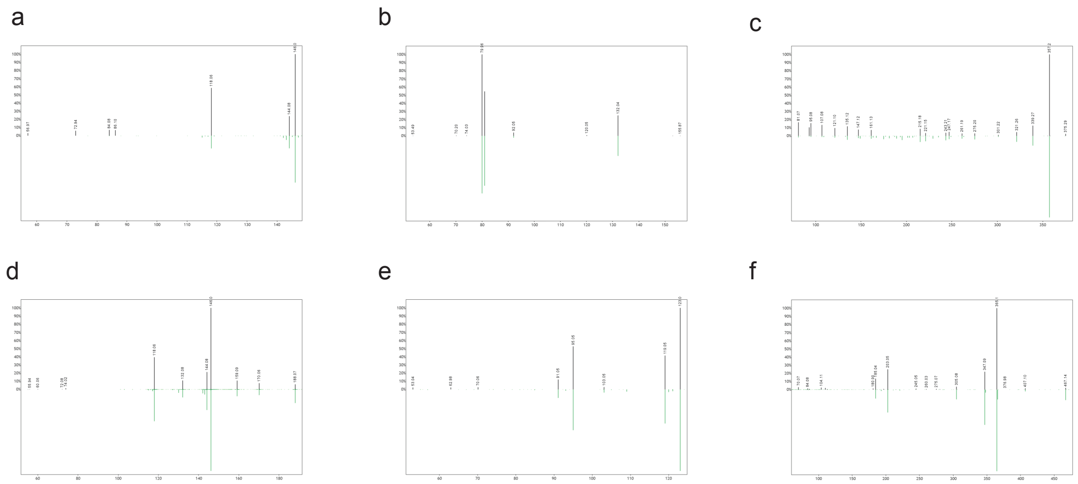

**Fig. S8. Representative microbially-modified, microbially-derived and microbially-influenced molecules detected in this dataset.** All annotations are level 2/3 according to the metabolomics standards initiative [36]. (a) *m/z* 188.071 RT 2.18 min (top, black) match to indole-l-lactate library reference (bottom, green). (b) *m/z* 212.002 RT 2.28 min (top, black) match to indoxyl sulfate library reference (bottom, green). (c) *m/z* 785.589 RT 2.891 min (top, black) match to deoxycholic acid library reference (bottom, green). (d) *m/z* 205.097 RT 1.24 min (top, black) match to tryptophan library reference (bottom, green). (e) *m/z* 165.055 RT 0.403 min (top, black) match to tyrosine library reference (bottom, green). (f) *m/z* 527.158 RT 0.265 min (top, black) match to maltotriose library reference (bottom, green).

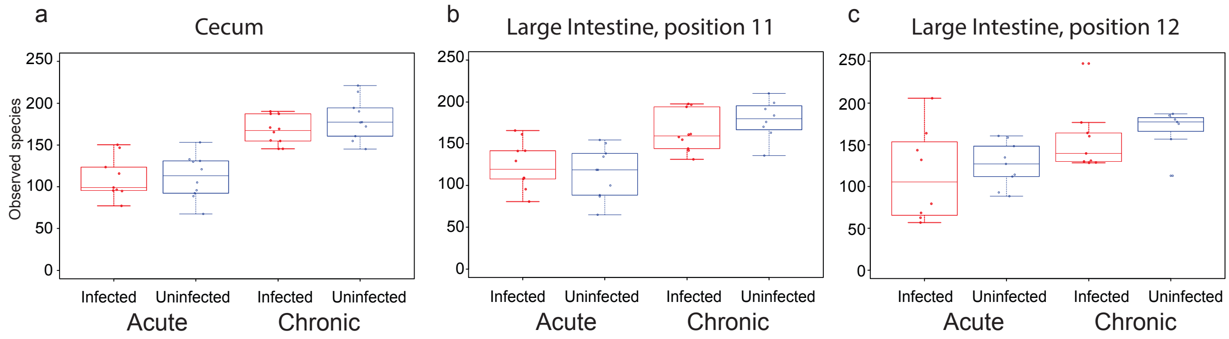

**Fig. S9. No impact of infection on alpha diversity.** (a) Cecum. (b) Proximal large intestine (sampling position 11). (c) Central large intestine (sampling position 12).

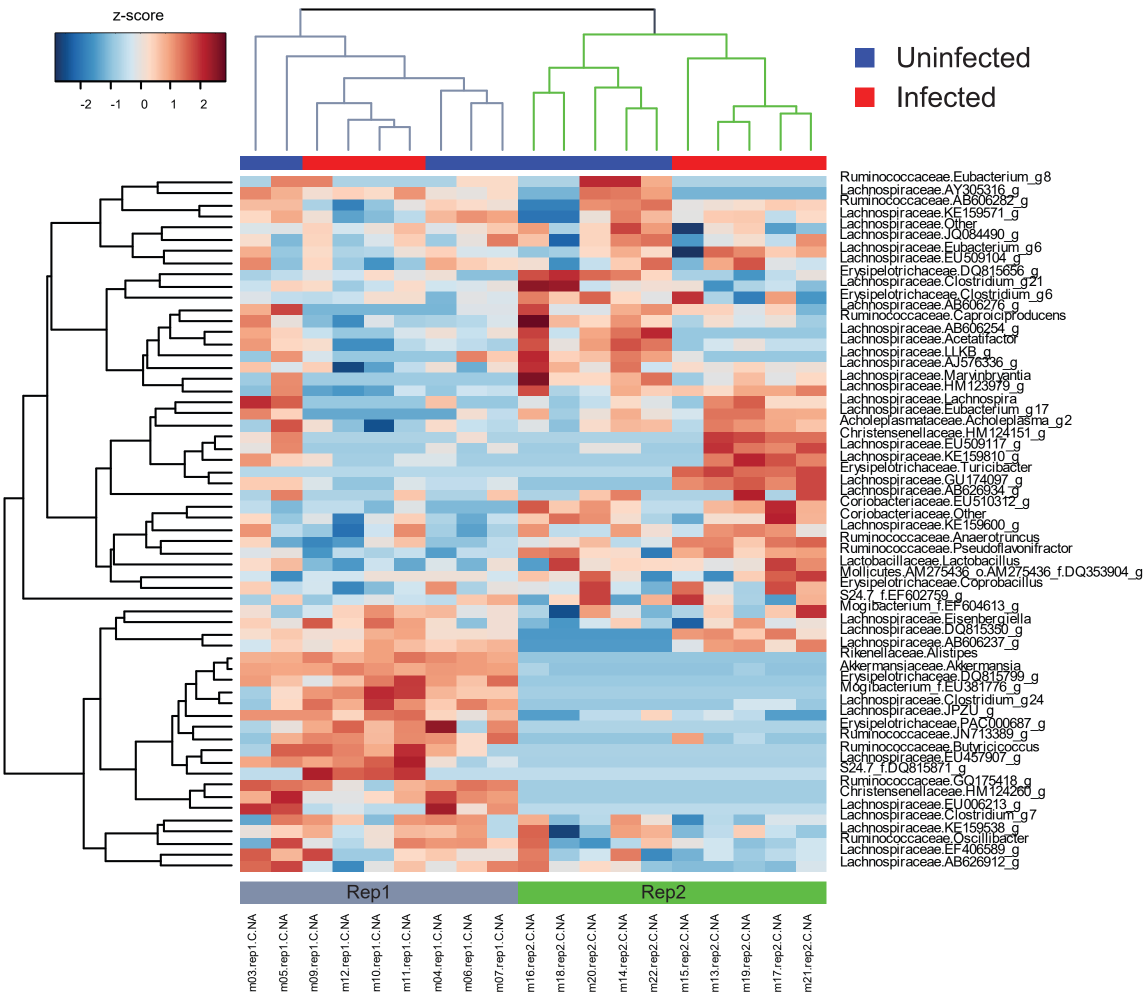

**Fig. S10. Batch effects in microbiome analysis.**

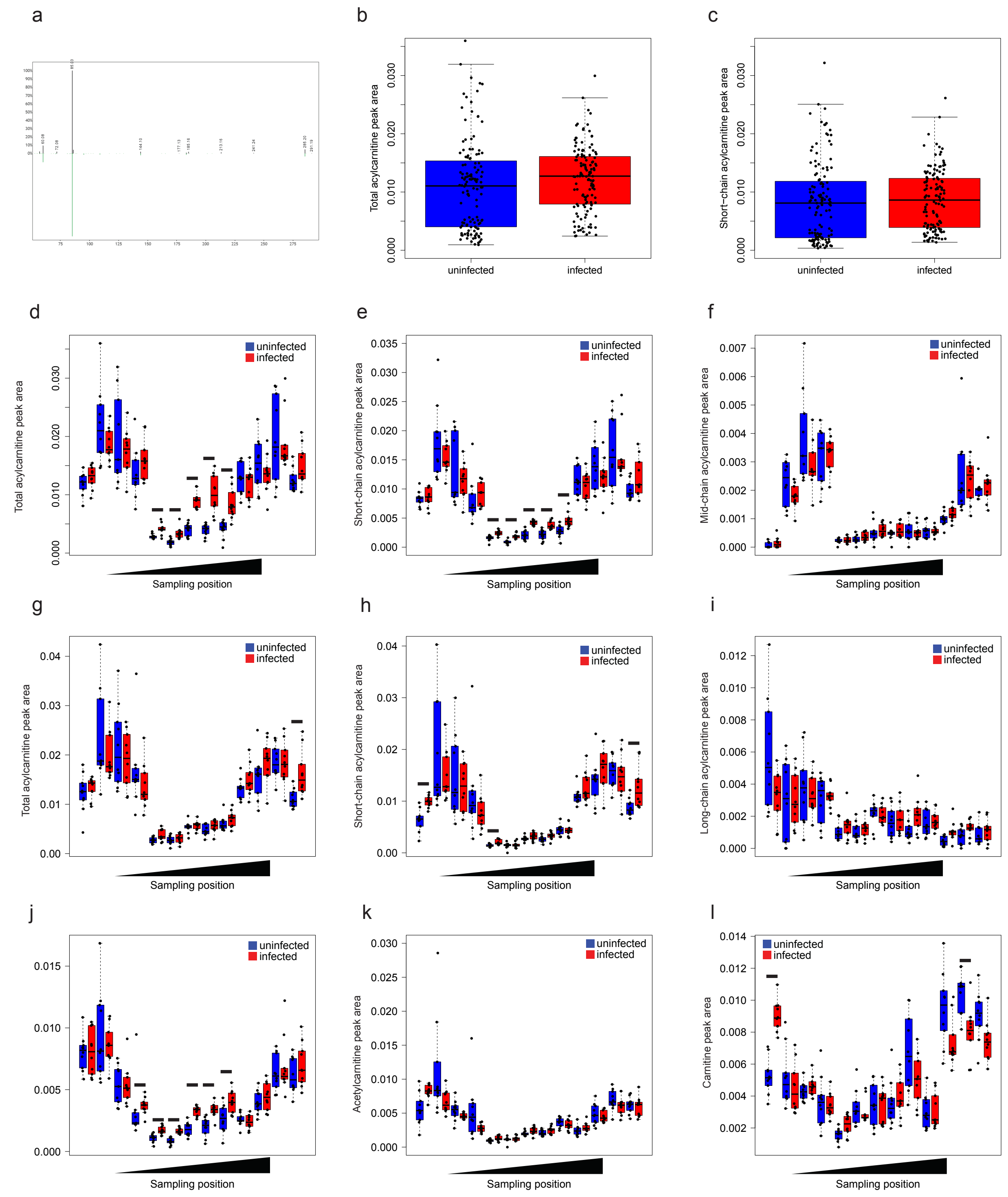

**Fig. S11. Carnitine, acetylcarnitine and acylcarnitine abundance.** (a) GNPS mirror plot showing MS2 match of *m/z* 344.279 RT 2.752 min (top, black) to lauroylcarnitine library reference (green, bottom). (b) Total acylcarnitine peak area across all GI sites, 12 days post-infection (p=0.01522 Mann-Whitney). (c) Total short-chain acylcarnitine peak area across all GI sites, 12 days post-infection (N/S Mann-Whitney). (d) Total acylcarnitine peak area at each sampling site, 12 days post-infection (FDR-corrected Mann-Whitney p=0.00543, p=0.000422, p=7.04e-05, p=7.04e-05, p=0.000422 for positions 5, 6, 7, 8, 9). (e) Short-chain acylcarnitine peak area at each sampling site, 12 days post-infection (FDR-corrected Mann-Whitney p=0.000668, p=0.000563, p=0.000141, p=0.000563, p=0.00189 for positions 5, 6, 7, 8, 9). (f) Mid-chain acylcarnitine peak area at each sampling site, 12 days post-infection. (g) Total acylcarnitine peak area at each sampling site, 89 days post-infection (FDR-corrected Mann-Whitney p=0.0196 for distal large intestine). (h) Short-chain acylcarnitine peak area at each sampling site, 89 days post-infection (FDR-corrected Mann-Whitney p=0.00422 for oesophagus; p=0.0253 for sampling position 5; p=0.0387 for distal large intestine). (i) Long-chain acylcarnitine peak area at each sampling site, 89 days post-infection. (j) Acetylcarnitine peak area at each sampling site, 12 days post-infection (FDR-corrected Mann-Whitney p=0.0.0220, p=0.003413, p=0.000141, p=0.000141, p=0.000891, p=0.0220 for sites number 4-9). (k) Acetylcarnitine peak area at each sampling site, 89 days post-infection. (l) Carnitine peak area at each sampling site, 89 days post-infection (FDR-corrected Mann-Whitney p=0.000141 for oesophagus and p=0. 0187 for central large intestine). Black bars indicate FDR-corrected Mann-Whitney p<0.05.

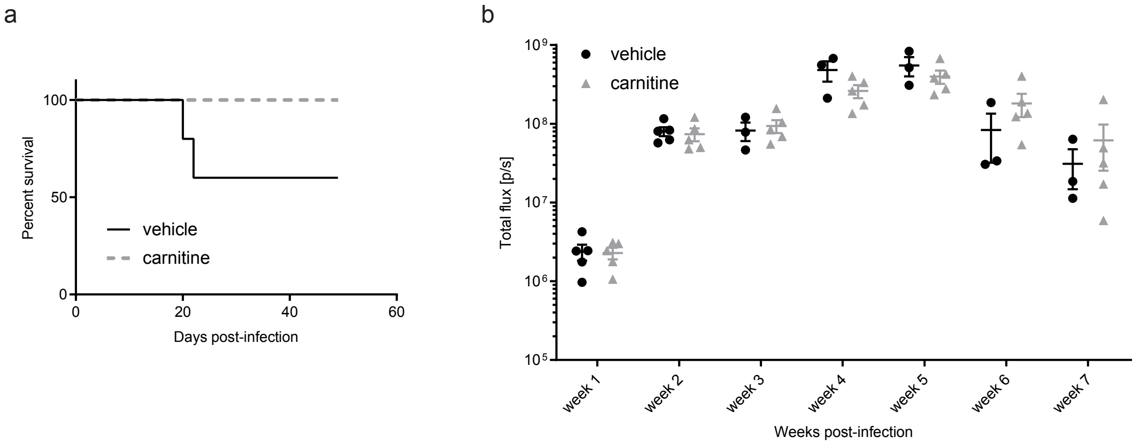

**Fig. S12. Impact of carnitine treatment on acute CD, intermediate-dose infection.** Male C3H/HeJ mice were infected with 5,000 luciferase-expressing *T. cruzi* strain CL Brener trypomastigotes. Beginning 7 days post-infection, animals received drinking water supplemented with 1.3% carnitine (100 mg/kg/day equivalent, based on water consumption) or remained on standard drinking water (vehicle group). (a) Acute-stage humane endpoints and mortality were only observed in the vehicle-treated group. (b) Comparable total body parasite burden between groups, weeks 1-7 post-infection. Mean + standard error of mean are displayed.
